# Old vs. New Local Ancestry Inference in HCHS/SOL: A Comparative Study

**DOI:** 10.1101/2025.02.04.636481

**Authors:** Xueying Chen, Hao Wang, Iris Broce, Anders Dale, Bing Yu, Laura Y Zhou, Xihao Li, Maria Argos, Martha L Daviglus, Jianwen Cai, Nora Franceschini, Tamar Sofer

**Affiliations:** Department of Biostatistics, Harvard T.H. Chan School of Public Health, Boston, MA, USA; CardioVascular Institute (CVI), Beth Israel Deaconess Medical Center, Boston, MA, USA; Department of Radiology, University of California San Diego, La Jolla, CA, USA; Department of Neurosciences, University of California, San Diego, San Diego, California, USA; Weill Institute for Neurosciences, Department of Neurology, University of California, San Francisco, UCSF, San Francisco, California, USA; Department of Epidemiology, School of Public Health, The University of Texas Health Science Center at Houston, Houston, TX, USA; Department of Biostatistics and Health Data Science, Indiana University School of Medicine, Indianapolis, IN, USA; Department of Biostatistics, University of North Carolina at Chapel Hill, Chapel Hill, NC, USA; Department of Genetics, University of North Carolina at Chapel Hill, Chapel Hill, NC, USA; Department of Environmental Health, School of Public Health, Boston University, Boston, MA, USA; Department of Epidemiology and Biostatistics, School of Public Health, University of Illinois Chicago, Chicago, IL, USA; Institute for Minority Health Research, University of Illinois at Chicago, Chicago, IL, USA; Department of Epidemiology, University of North Carolina at Chapel Hill, Chapel Hill, NC, USA; Department of Medicine, Harvard Medical School, Boston, MA, USA; Department of Medicine, Brigham and Women’s Hospital, Boston, MA, USA

## Abstract

Hispanic/Latino populations are admixed, with genetic contributions from multiple ancestral populations. Studies of genetic association in these admixed populations often use methods such as admixture mapping, which relies on inferred counts of “local” ancestry, i.e., of the source ancestral population at a locus. Local ancestries are inferred using external reference panels that represent ancestral populations, making the choice of inference method and reference panel critical. This study used a dataset of Hispanic/Latino individuals from the Hispanic Community Health Study/Study of Latinos (HCHS/SOL) to evaluate the “old” local ancestry inference performed using the state-of-the-art inference method, RFMix, alongside “new” inferences performed using Fast Local Ancestry Estimation (FLARE), which also used an updated reference panel. We compared their performance in terms of global and local ancestry correlations, as well as admixture mapping-based associations. Overall, the old RFMix and new FLARE inferences were highly similar for both global and local ancestries, with FLARE-inferred datasets yielding admixture mapping results consistent with those computed from RFMix. However, in some genomic regions the old and new local ancestries have relatively lower correlations (Pearson R < 0.9). Most of these genomic regions (86.42%) were mapped to either ENCODE blacklist regions, or to gene clusters, compared to 7.67% of randomly-matched regions with high correlations (Pearson R > 0.97) between old and new local ancestries.

## Introduction

Admixture is the process where “ancestral” populations that have been isolated interbreed to form a new, admixed population, where specific segments in genomes of individuals from the admixed population can be traced to one of the ancestral populations[1]. Admixed individuals with multiple ancestral populations have diverse genetic compositions leading to differential linkage disequilibrium patterns both across ancestral and within admixed populations[2]. Further, disease risk or quantitative trait loci often have different frequencies across ancestral populations. Thus, inference of genetic architecture of complex traits in admixed populations is complicated and more challenging compared to other well-studied populations that are genetically more homogeneous [3]. Hispanic/Latino populations are admixed, and their admixture patterns are typically described based on European, African, and Amerindian continental ancestral populations [4,5]. Hispanic/Latino individuals account for 17.8% of the United States’ work-age population in 2023 as the largest racial/ethnic minority group [6]. However, the group is still underrepresented in genomic studies. According to the genome-wide association studies (GWAS) Diversity Monitor, accessed in July 2024, only 0.38% of GWAS participants were identified as Hispanic or Latin American [7]. Identification of causal variants and of their effect magnitudes in Hispanic/Latino individuals is thus complicated by the limited representation and complexities from their admixture. Overcoming these methodological challenges and efficiently detecting potential variants and effects remain as knowledge gaps to be addressed.

Nevertheless, prior research on admixed populations has effectively utilized admixture information from ancestral populations, defined according to external reference panels, to identify disease-associated variants, and a few approaches have also been proposed to leverage local ancestries for genetic association analysis with a phenotype or trait [4]. For instance, the two-step testing procedure LAAA (Local Ancestry Adjusted Allelic) uses an omnibus joint test to examine the effects in allele, local ancestry, and ancestry plus alleles, and then uses model selection to detect the sources of associations [8]. The TRACTOR method performs ancestry-specific GWAS with a regression model that is local ancestry-aware to generate effect sizes and p-value for each ancestry [9]. Genetic analyses, such as GAUDI, incorporate local ancestries into polygenic risk scores, which model ancestry-differential effects using penalized regression frameworks [10].

Amidst these methodologies, admixture mapping emerges as an effective analytic approach for analyzing association with a phenotype using local ancestry [11–13]. Local ancestry is defined at the local, i.e., variant or genomic segment level, referring to the reference ancestry from which the locus was inherited. In contrast, global ancestry is defined based on patterns across the genome: it estimates the proportion of genome inherited from ancestors from a specified (or inferred) genetic ancestry, i.e., it is an average of local ancestry patterns across the genome. At a given genomic position, the number of chromosomal copies with local ancestry from a given ancestry is coded (possibly 0, 1, or 2 copies) [14], and this variable is used in association analysis with the phenotype. An association is observed when either the causal variant frequency or effect sizes (or both) differ between the modeled ancestral populations [15]. The modeled local ancestry unit (variant or segment) typically captures the causal variant via linkage disequilibrium, where the range of LD is usually very high for local ancestry [16]. As a result, a phenotype association with local ancestry may be driven by genetic variants located at a potentially large genomic segment around the region of association [17]. Admixture mapping is powerful because it may capture more complex associations (haplotypes, rare variants) compared to standard GWAS and has a lower multiple testing burden [13,18]. Admixture mapping can also be useful to augment standard variant-level analysis now that more modern and computationally efficient methods, as well as more genomic reference panels, are available to perform local ancestry inference [19–26]. For example, our team recently reported associations of local ancestries with blood metabolite levels in Hispanic Community Health Study/Study of Latinos (HCHS/SOL) participants, yielding 116 novel associations with 78 circulating metabolites, in which genomic regions enriched for both African ancestry and Amerindian ancestry were identified respectively and mapped to the corresponding genes with their associated metabolic pathways and diseases [27].

Local ancestry can be inferred through reference panels with different ancestral populations. Methods include a discriminative model trained on reference panels (RFMix) and generative models, such as MOSAIC and HapMix [14,28,29]. In the HCHS/SOL, over ten thousand study participants were genotyped, and their local ancestries were inferred [5,30,31]. HCHS/SOL local ancestry inference was reported in previous works, where the inference of local ancestries was described over genomic intervals [5]. This published study used RFMix, which was considered state-of-the-art at that time [29]. The local ancestry inference accounted for three ancestral, continental populations: Europe, Africa, and America (a few individuals with non-negligible contributions of Asian ancestry were identified and excluded from analysis). From this inference, numerous publications reported discoveries of genetic associations via admixture mapping followed by overlaying of GWAS results on the admixture mapping signal [15,27,32,33]. More recently, FLARE (Fast Local Ancestry Estimation), a new method for local ancestry inference, reported improved performance at both the computational and inference accuracy level compared to past approaches using both simulated and real data from 1000 Genomes and Human Genome Diversity Project [19,20,34]. Thus, FLARE can be applied to the HCHS/SOL participants, generating detailed inferences referencing either the previous three populations or by including additional reference ancestral populations, thanks to its enhanced computational capacity. This study quantifies the potential improvements of the FLARE method through comprehensive comparisons of local ancestry inference and subsequent admixture mapping between the RFMix and the FLARE inferences.

This study has two goals. First, it aims to compare the resulting local ancestry estimates from FLARE to those of the RFMix inference. For the FLARE inference, the analyses involved applying FLARE with two sets of reference populations: once with 7 continental ancestries (FLARE7) and once with the 3 continental ancestries previously used (FLARE3). Second, it compares the use of these inferred local ancestries in admixture mapping analysis applied to metabolites that were previously reported to be associated with local ancestry levels in specific genomic regions [27].

## Results

### Participant characteristics

Participant characteristics focused on individuals participating in the metabolomics analysis and are provided in Table 1, stratified by metabolomics batch. The mean age was ∼46 in the discovery batch (batch 1), and ∼52 in the replication batch (batch 2). Batch 2 also had a larger proportion of female participants (∼64% compared with ∼57%), higher proportion of participants with hypertension and/or diabetes, and lower eGFR levels on average. For both batches, the distribution of participants from the various recruitment centers were similar (Table 1).

**Table 1.** Characteristics of participants included in the admixture mapping, from either the discovery batch 1 or the replication batch 2. Continuous variables are reported with mean and standard deviation (SD) and categorical variables are reported with proportion in each level.

|  | Discovery Batch<br>(N=3861) | Replication Batch<br>(N=1644) | Overall<br>(N=5505) |
| --- | --- | --- | --- |
| **Age (years)** |  |  |  |
| Mean (SD) | 45.9 (13.8) | 52.3 (11.0) | 47.8 (13.4) |
| **BMI (kg/m<sup>2</sup>)** |  |  |  |
| Mean (SD) | 29.8 (6.07) | 30.2 (6.06) | 29.9 (6.07) |
| **eGFR (mL/min/1.73m<sup>2</sup>)** |  |  |  |
| Mean (SD) | 108 (26.6) | 106 (26.6) | 108 (26.6) |
| **Gender** |  |  |  |
| Female | 2217 (57.4%) | 1045 (63.6%) | 3262 (59.3%) |
| Male | 1644 (42.6%) | 599 (36.4%) | 2243 (40.7%) |
| **Recruitment Center** |  |  |  |
| Bronx | 1025 (26.5%) | 442 (26.9%) | 1467 (26.6%) |
| Chicago | 884 (22.9%) | 368 (22.4%) | 1252 (22.7%) |
| Miami | 1078 (27.9%) | 514 (31.3%) | 1592 (28.9%) |
| San Diego | 874 (22.6%) | 320 (19.5%) | 1194 (21.7%) |
| **Genetic Analysis Group** |  |  |  |
| CentralAmerican | 387 (10.0%) | 182 (11.1%) | 569 (10.3%) |
| Cuban | 735 (19.0%) | 357 (21.7%) | 1092 (19.8%) |
| Dominican | 377 (9.8%) | 210 (12.8%) | 587 (10.7%) |
| Mexican | 1397 (36.2%) | 481 (29.3%) | 1878 (34.1%) |
| PuertoRican | 707 (18.3%) | 277 (16.8%) | 984 (17.9%) |
| SouthAmerican | 258 (6.7%) | 137 (8.3%) | 395 (7.2%) |
| **Hypertension** | 1104 (28.6%) | 650 (39.5%) | 1754 (31.9%) |
| **Diabetes** | 734 (19.0%) | 476 (29.0%) | 1210 (22.0%) |

### Global ancestry comparison

The number of variants that passed this quality control (QC) is 3,610,937 single nucleotide polymorphisms (SNPs) for FLARE3 (99.78%) and 5,091,756 SNPs for FLARE7 (99.74%). The global ancestry proportions for each genetic analysis group are highly similar between RFMix and FLARE3 results (Supp. Figure 1A, 1B). Inference of European, African, and Amerindian ancestry reiterate previously reported findings for this datasets [13,27,35]: participants from the Cuban group have more European ancestry than other groups, and individuals from Mainland backgrounds (Mexico, Central America, and South America) have more Amerindian ancestry than Caribbean groups. By contrast, participants from Caribbean backgrounds (Dominican, Puerto Rican) have more African ancestry compared to the participants from mainland backgrounds. The introduction of multiple other ancestries in the FLARE7 inference added more details to the ancestry compositions (Supp. Figure 1C). The estimated global proportions for the four additional ancestries (Central/South Asia, East Asia, Oceania, and Middle East) for most individuals are less than 5%. No participants have Oceanian or Central/South Asian ancestries. The Cuban group has more Middle Eastern ancestry than other groups. The groups with Caribbean backgrounds have the least East Asian ancestry (Supp. Figure 1C).

Comparing the estimated global ancestry proportions in the two FLARE inferences and RFMix to those from ADMIXTURE, we see that all correlations are above 0.99, suggesting high similarity (Figure 1). This can also be observed from visualization of the first two genetic PCs colored by the global ancestry proportions computed from RFMix, FLARE3, and FLARE7 (Supp. Figure 2). As previously reported[5], RFMix tends to have higher European ancestry calls than ADMIXTURE, which assigns lower European ancestry and higher African and Amerindian ancestry proportions. A similar pattern is also observed in local ancestry calls when comparing FLARE3 to ADMIXTURE proportions (Figure 1). In FLARE7, the European ancestry proportions tend to be lower for some individuals compared to those in RFMix, ADMIXTURE, and FLARE3, as local ancestry counts could be attributed to other ancestries such as Middle Eastern or East Asian (Figure 1, Supp. Figure 1C).

**Figure 1.**
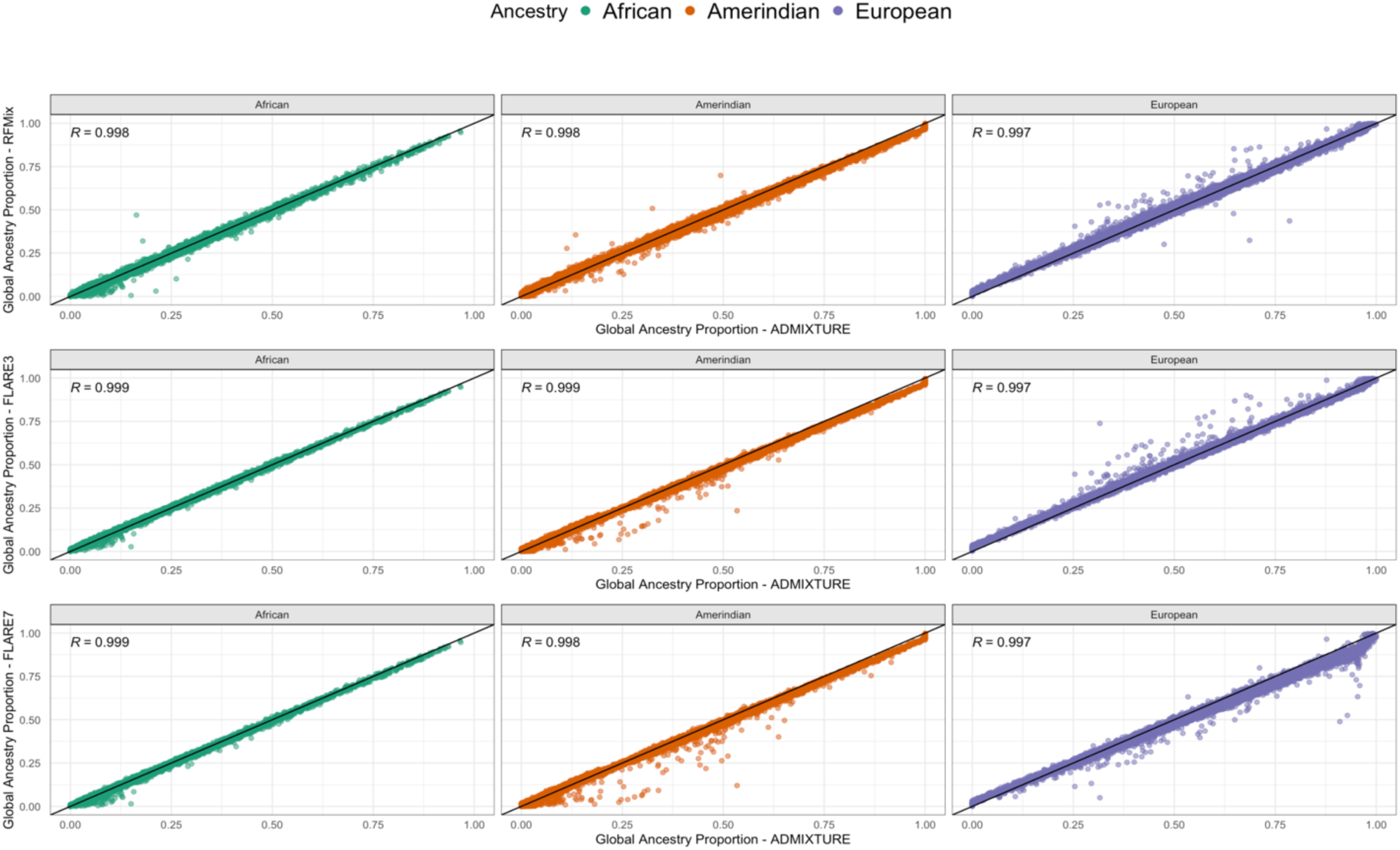
Global ancestry correlation comparisons for RFMix, FLARE3, and FLARE7 to ADMIXTURE. Estimated proportions of global ancestries (African, Amerindian, and European) from two inferences are plotted against each other, and the Pearson correlation coefficient (R) is provided for each comparison.

### Local ancestry comparison

For the entire genome, 15425 (99.52%) local ancestry intervals from the RFMix-based dataset were successfully converted into the hg38 assembly. 92.29% of the SNPs from FLARE7 and 92.48% of the SNPs from FLARE3 were matched to an RFMix-inferred local ancestry interval. For all ancestries, the mean correlations between the RFMix and FLARE3 local ancestry counts are above 0.9, with values of 0.954 (standard deviation [SD] = 0.012) for African, 0.972 (SD = 0.009) for Amerindian, and 0.956 (SD = 0.011) for European ancestry. The mean correlations between RFMix and FLARE7 are lower for European ancestry and comparable for the other two ancestries, with values of 0.959 (SD = 0.012) for African, 0.972 (SD = 0.009) for Amerindian, and 0.916 (SD = 0.013) for European ancestry. Compared to FLARE3, European ancestry in FLARE7 has lower correlations with the European ancestry counts in the RFMix dataset. This could be due to the introduction of Middle Eastern ancestry, where some parts of the local ancestry previously assigned to European are now assigned to Middle Eastern ancestry (Figure 2, Supp. Figure 1C). These differences are also reflected in the comparison between mean proportions of European ancestry for each SNP position. The RFMix and FLARE3 inferences consistently have higher European proportions than those of FLARE7 across all chromosomes (Supp. Figure 4, 5). The variances of the estimated ancestry proportions computed across the genome in both FLARE3 and FLARE7 are highly similar and are higher than that of RFMix across ancestries and chromosomes (Supp. Figure 6, 7).

**Figure 2.**
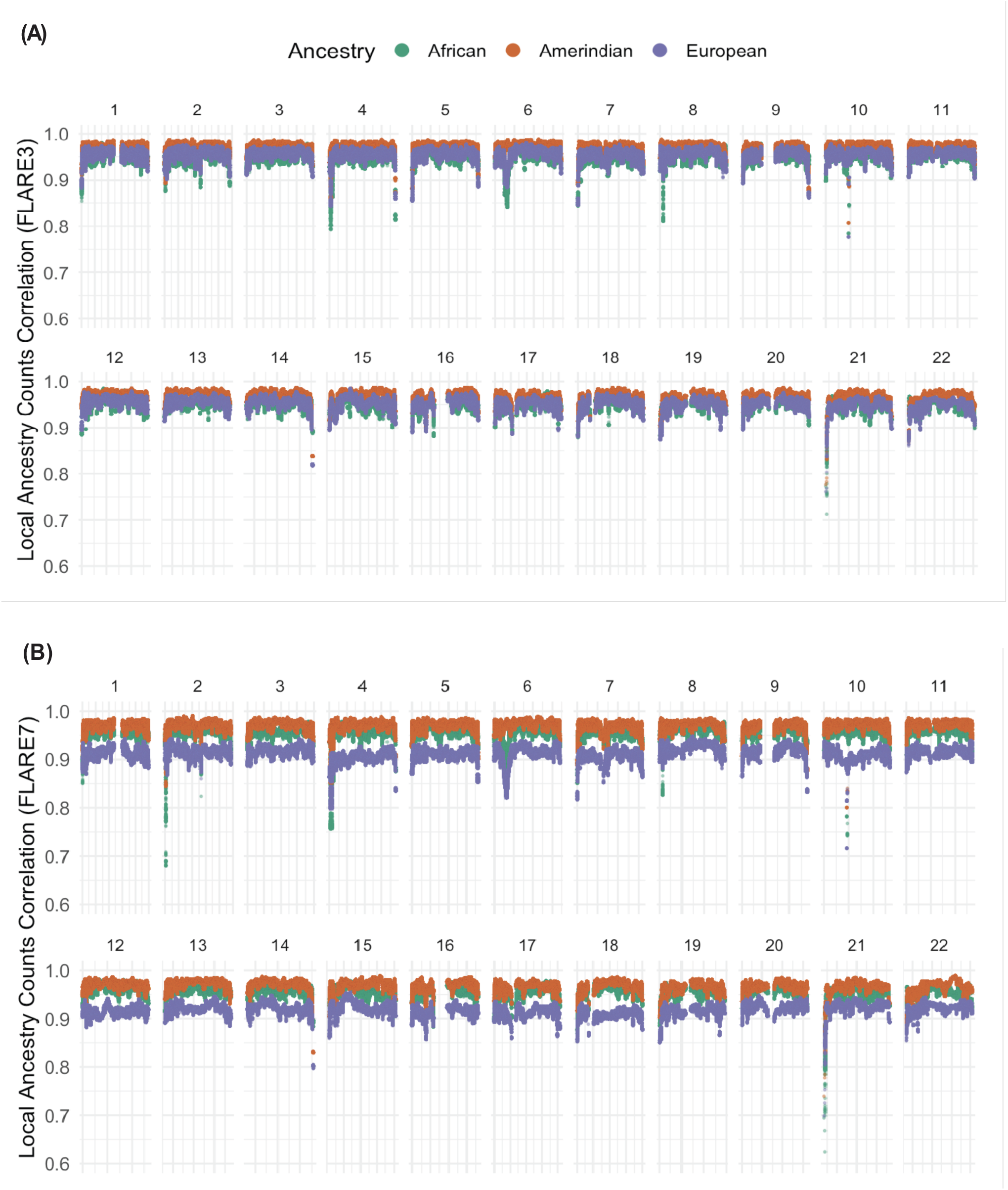
Pearson correlation between the RFMix and FLARE3 (A) and between RFMix and FLARE7 (B) for each matched SNP and ancestry block across chromosomes 1 to 22.

For certain chromosomes there are regions where the correlations between the RFMix and the FLARE inferences drop at the start or/and end position of a chromosome, mostly observed in chromosome 2, 4 and 21 (Figure 2). There are less correlation drops in FLARE3 vs. RFMix than that in FLARE7 vs. RFMix. This inconsistency between inferences reflects differences in inferred mean proportions of different ancestries. In chromosome 2, the mean African proportion decreased, relative to the RFMix inference at the starting regions, whereas the Amerindian ancestry proportion increased (Supp. Figure 4, 5). However, in chromosomes 10 and 21, there are regions where both the mean Amerindian and European ancestry proportions dropped while the African ancestry proportion increased (Supp. Figure 4, 5). A similar pattern is observed in variances of African and Amerindian ancestry counts (Supp. Figure 6, 7).

We also considered a stricter QC metric, increasing the imputation quality threshold R^2^ applied on variants from the FLARE datasets from 0.8 to 0.95, to assess its impact on the low-correlation regions between RFMix and FLARE3 inferences. Even after filtering, the SNP (FLARE3) – interval (RFMix) pairs continued to exhibit low correlations in the same regions (Supp. Figure 3). These low-correlation regions were subsequently examined in greater detail. In the comparison of FLARE3 and RFMix inferences, 31,148 pairs (0.93%) showed low correlation (correlation coefficient < 0.9) in any ancestry. Of these, 26,921 pairs (86.42%) were mapped to either ENCODE blacklist (10,340 pairs, 38.41%) or annotated gene clusters (18,399 pairs, 68.34%) in UCSC genome browser, overall corresponding to 54 unique annotated genomic regions (Supp. Table 1). By contrast, for the high-correlation pairs (correlation coefficient > 0.97) that were randomly selected, 3350 (7.67%) were mapped to the ENCODE blacklist and none were in proximity to the identified gene clusters. This suggests that the low correlations are enriched in the mapped genomic regions.

### Admixture mapping results

The most statistically significant chromosomal regions from the admixture mapping and the ancestry driving the phenotypic differences align with the published findings: N-acetylarginine (chromosome 2, driven by African ancestry), 3-aminoisobutyrate (chromosome 5, driven by Amerindian ancestry), PC 16:0/20:4 (chromosome 11, driven by Amerindian ancestry), and PE 16:0/20:4 (chromosome 15, driven by Amerindian ancestry) (Table 2). The computed significance thresholds of batch 1 from the RFMix (2.05 × 10^-6^), FLARE3 (2.58 × 10^-6^), and FLARE7 (1.95 × 10^-6^) are similar. After accounting for genome-wide multiple testing burden in each inference, the test statistics in both African and Amerindian ancestries show statistically significant associations at the same chromosomal regions for PC 16:0/20:4 (Table 2). However, the association is stronger for Amerindian ancestry, suggesting that the unknown underlying causal variant (or variants) have stronger frequency difference when comparing Amerindian to combined European and African local ancestries, than the difference when comparing African ancestry to the combined European and Amerindian local ancestries.

**Table 2.** Comparison of p-values from the most significant variant associations (p*) detected in the most statistically significant associated genomic regions from RFMix admixture mapping results. The p* values are compared across the three inferences (RFMix, FLARE3, and FLARE7) as well as between the discovery batch 1 (b1) and replication batch 2 (b2).

| Metabolite | Ancestry | Region(RFMix) | p* RFMix b1 | p* RFMix b2 | rsID FLARE3 | p* FLARE3 b1 | p* FLARE3 b2 | rsID FLARE7 | p* FLARE7 b1 | p* FLARE7 b2 |
| --- | --- | --- | --- | --- | --- | --- | --- | --- | --- | --- |
| **N-acetylarginine** | African | chr2(p13.2) | 1.603e-10 | 1.409e-07 | rs56016348 | 8.188e-10 | 8.722e-09 | rs75054060 | 3.975e-10 | 7.473e-07 |
| **3-aminoisobutyrate** | Amerindian | chr5(p13.2) | 1.928e-35 | 2.632e-22 | rs371913 | 4.090e-35 | 4.384e-22 | rs10941230 | 6.427e-37 | 3.551e-23 |
| **PC 16:0/20:4** | African | chr11(q12.2) | 3.251e-10 | 2.287e-06 | rs112534078 | 1.169e-09 | 2.300e-05 | rs55970610 | 3.241e-10 | 8.697e-06 |
| **PC 16:0/20:4** | Amerindian | chr11(q12.2) | 1.675e-50 | 5.828e-26 | rs174573 | 3.175e-50 | 9.983e-25 | rs7935946 | 2.002e-50 | 9.639e-24 |
| **PE 16:0/20:4** | Amerindian | chr15(q21.3) | 2.776e-12 | 2.766e-05 | rs8192701 | 1.822e-13 | 1.195e-05 | rs74248033 | 1.112e-12 | 9.488e-05 |

Within each batch and metabolite, the p-value levels for the most significantly associated genomic region (p*) identified using RFMix local ancestry inference are very similar to the p-values of the most significant variants estimated from both FLARE3 and FLARE7 at the same region (Table 2, Figure 3, Figure 4). In the most statistically significant regions, most of the identified variants are in high proximity to the associated genes previously reported (Supp. Figure 9, 11, 12). Several of the most significant variants are located at a greater distance from the reported genes (Supp. Figures 8, 10). Notably, this includes variants identified through admixture mapping with N-acetylarginine on chromosome 2 and a variant associated with PC 16:0/20:4 via FLARE7 inference in batch 1 individuals using African local ancestry. Nonetheless, all identified variants are situated within 1 Mb of the corresponding reported genes.

**Figure 3.**
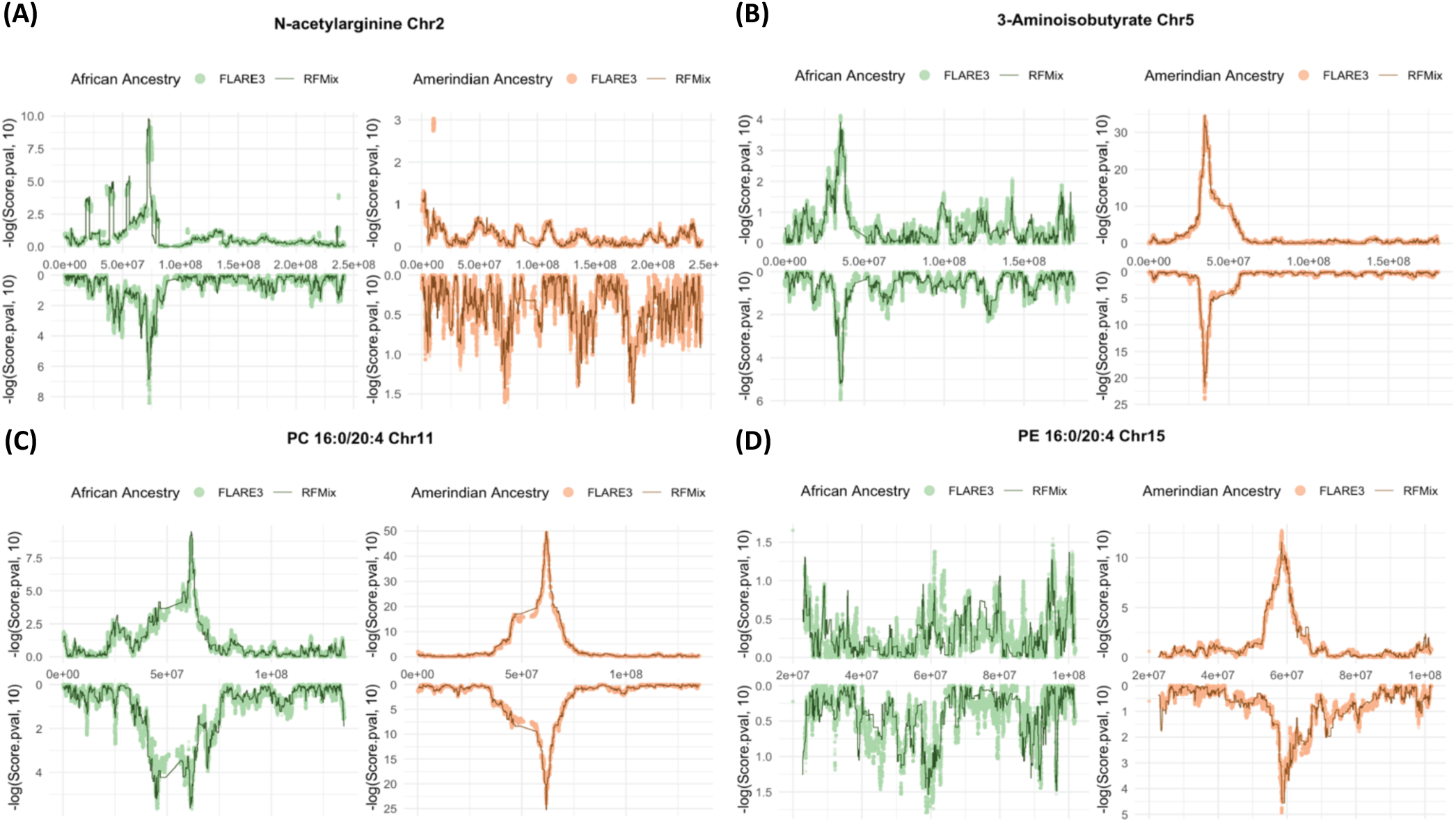
Admixture mapping using local ancestry counts in RFMix and FLARE3 at the chromosome with most significant admixture mapping associations using the driving local ancestry for the four metabolites: N-acetylarginine at chromosome 2 (A), 3-aminoisobutyrate at chromosome 5 (B), PC 16:0/20:4 at chromosome 11 (C), and PE 16:0/20:4 at chromosome 15 (D). Results from the discovery and replication datasets are displayed as mirrored plots, with the upper panel representing the discovery (batch 1) and the lower panel representing the replication (batch 2) datasets.

**Figure 4.**
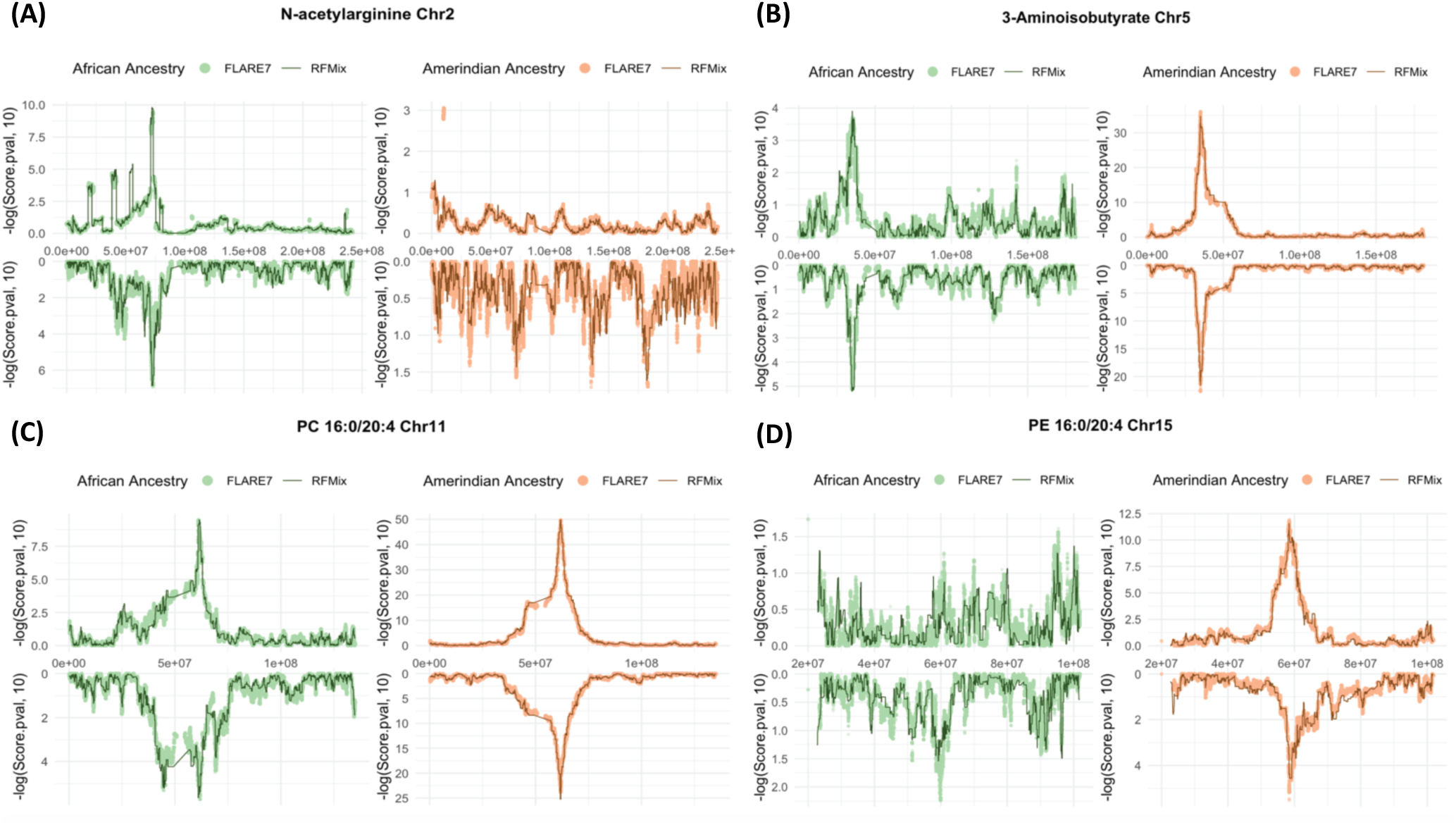
Admixture mapping using local ancestry counts in RFMix and FLARE7 at the chromosome with most significant admixture mapping associations using the driving local ancestry for the four metabolites: N-acetylarginine at chromosome 2 (A), 3-aminoisobutyrate at chromosome 5 (B), PC 16:0/20:4 at chromosome 11 (C), and PE 16:0/20:4 at chromosome 15 (D). Results from the discovery and replication datasets are displayed as mirrored plots, with the upper panel representing the discovery (batch 1) and the lower panel representing the replication (batch 2) datasets.

Among the regions inside the relatively low correlations between RFMix and FLARE inferences (< 0.9), in any of the ancestries, interval chr16:29,222,050-29,296,362 is mapped to the ENCODE blacklist and the region has a statistically significant association to propyl 4-hydroxybenzoate sulfate previously reported (Supp. Table 1) [27]. The local African ancestry association in this low correlation region is less significant in FLARE datasets than RFMix (Supp. Figure 13). Besides this region of low correlation in local ancestry counts, admixture mapping results from both FLARE inferences identify a significant association region with African Ancestry beginning from 16q11.2, where in RFMix there were no mapped intervals so that the association region begins at 16q12.1 (Supp. Figure 13). This may be due to the RFMix inference being performed using the hg19 coordinates.

## Discussion

Past local ancestry inference in HCHS/SOL used RFMix and was based on a limited number of genotypes. It was the basis for multiple admixture mapping analyses, and we expect the uses of local ancestry inference to continue to develop. As such, it is important to compare the past inference to that by FLARE – a new popular method that is being applied to HCHS/SOL and other datasets. We compared the RFMix and new FLARE3 and FLARE7 inferences, that rely on 3 and 7 reference populations respectively, via visualizations and correlation in global ancestry proportions, local ancestry counts across the genome, and admixture mapping focusing on a few previously reported association regions.

Estimated global ancestry proportions of Amerindian, African, and European ancestries were similar between the three inferences, and similar to global ancestry proportions previously computed using ADMIXTURE. The few observed discrepancies between RFMix, FLARE3, and ADMIXTURE may arise because ADMIXTURE models ancestry based on allele frequencies across the genome [5,36]. This results in some assignments of portions of European ancestry to African and Amerindian components, resulting in deflated European ancestry estimates and inflated estimates for the other ancestries compared, in comparison to inferences based on local analyses (Figure 1). Local proportions computed over HCHS/SOL individuals across the genome provide a more detailed view, demonstrating that local ancestry counts between RFMix and both FLARE inferences are highly correlated, as suggested in the previous study using real data from admixed individuals [37]. Additionally, as expected, that the correlations between RFMix and FLARE7 ancestry counts are lower compared to those between RFMix and FLARE3, due to the use of a different set of reference populations. The correlations between RFMix and FLARE7 European ancestry counts dropped (compared to RFMix-FLARE3 comparison) across the genome, from an average of 0.956 to an average of 0.916, likely because some genomic segments that have been attributed to European ancestry in FLARE3 were assigned to Middle Eastern ancestry in the FLARE7 inference. There are patterns of fluctuating correlations between RFMix and FLARE local ancestries across the genome for both FLARE3 and FLARE7 (Figure 2). Some variability is expected, especially as FLARE inferences relied on a denser set of SNPs. Specific regions showed substantial differences in correlations, with observed correlation lower than 0.9. In the most extreme case, the correlations between RFMix and FLARE7 local African ancestries were less than 0.7. A few notable regions where correlations between local ancestry counts dropped substantially compared to the genome-wide average are close to the telomeres, in proximity to either the regions from the ENCODE blacklist or gene clusters annotated by UCSC genome browser (Supp. Table 1). For example, a region with a drop in correlation between RFMix and both FLARE local ancestry inferences at chromosome 6 is around the major histocompatibility complex (MHC) region (chr6: 28,510,019-33,480,577), which is famous for containing clusters of closely linked supergenes that are related to immune responses [38,39]. The low correlation of local ancestry counts in the MHC region can be attributed to differences in the reference populations used. Both RFMix and FLARE infer an excess of African ancestry in the MHC region compared to genome-wide proportions (Supp. Figures 4, 5), which aligns with findings from multiple previous studies that reported a similar pattern of MHC-specific excess of African ancestry across various Latino/Hispanic populations[40–43]. However, the inference quality and potential selection mechanism in this region are still not well-understood due to the complexity of long-range linkage disequilibrium in the MHC region [40,44]. Another study on the local ancestry of admixed American individuals identified an excess of Amerindian ancestry on chromosome 8p23.1, an inverted region with extended LD, as well as a positive selection signal in this region for admixed populations but no selection signal when all Amerindian reference populations were pooled. Our analysis did not observe a significant excess of Amerindian ancestry at chromosome 8p23.1 (Supp. Figures 4, 5), which may be explained by the use of reference panels from multiple American populations. Note that a mapped blacklisted region for low local ancestry correlations is also located at chromosome 8p23.1 (Supp. Table 1), close to the reported inverted region [45]. It is possible that the FLARE inferences have better accuracy in these specific regions comparing to RFMix (and this could be related to differences in reference panel individuals and genotyping methods), but we cannot conclude that with certainty. The differences in local ancestries in genetically “complicated” regions, (blacklisted regions or gene clusters), which likely indicate higher variance and less certainty in local ancestry inference, do not appear to substantially affect the reported metabolite associations (Supp. Figure 14). However, we were unable to determine how these differences influenced the accuracy of the inference based on the observed variations in the correlations and the admixture mapping results.

FLARE7 includes more ancestral populations. However, the inclusion of multiple additional ancestries appears to have a limited practical impact. Participants did not have local ancestry counts identified as Central/South Asian or Oceanian, and most participants only had minimal East Asian ancestry. Consequently, these additional ancestry counts offer limited information and increase the computational burden. For Hispanic/Latino populations in the U.S., it is probably sufficient to use three continental ancestral populations (European, African, and Amerindian). Researchers may want to include more populations for compatibility with other datasets, for example.

For both FLARE inferences, the admixture mapping results for the four metabolites align with the previous admixture mapping associations identified using the RFMix dataset. The FLARE inferences provided a higher resolution to the variant level compared to that in the RFMix results, and their reproducibility across batches for the admixture mapping result is also verified[27] (Figure 2, 3). When comparing global and local ancestry and the admixture mapping results, note that the discrepancies can stem from multiple differences between datasets aside from the method of local ancestry inference: arrays and reference panels used in genotyping and genotype imputation, as well as the fact that the two FLARE datasets are based on imputed data from HCHS/SOL.

Because of the smaller number of individuals, a higher p-value (lower significance) from the results in the replication batch is expected (Table 2). As previously reported, metabolites in N-acetyl amino acid and is associated with a region in gene ALMS1, which was found to be associated with chronic kidney disease [27,46]. The SNP with the most significant association identified is rs56016348 and rs75054060 for batch 1 in FLARE3 and FLARE7, respectively, both of which are situated within 1Mb from the 5’ end of ALMS1 in terms of genetic distance (Supp. Figure 8).

For the association with 3-aminoisobutryate (beta-aminoisobutyric acid), the variants with the most statistically significant associations in the region for both FLARE3 and FLARE7 are clearly localized to the gene AGXT2, which was shown in HCHS/SOL to be associated with Mild Cognitive Impairment (MCI) in Hispanic/Latino adults, and the genetic association is mediated by the levels of beta-aminoisobutyric acid [47] (Supp. Figure 9). For PE:16:0/20:4 and PC:16:0/20:4, the variants with most statistically significant associations in both FLARE3 and FLARE7 are not in the exact same gene as previously identified variants (rs2070895 in LIPC and rs174562 in FADS2) [27], but still in high proximity with the reported genes (Supp. Figure 10, 11, 12). This should be interpreted with caution, because LD between local ancestries are extensive (long ranged) reducing precision in localizing signals; the observed statistically significant variants could be in high local ancestry LD with the identified variants that were found to explain the significant admixture mapping signals[17].

Despite the FLARE inference method enhancing the resolution of local ancestry information to the variant level, the presence of high LD still hampers identification of causal variants driving the admixture mapping signal. Thus, there is still a need of fine mapping using variant-level genotyping data[15,27].

In summary, we compared the old HCHS/SOL RFMix-based local ancestry inference to newer FLARE3 and FLARE7 inferences. The resulting local inferred European, African, and Amerindian ancestries are overall similar, with some differences that are clearly attributable to different reference populations (FLARE7 used 7 continental reference populations), and those that are possibly found in regions that are more difficult for genotyping (gene heavy, high LD) and perhaps for local ancestry inference as well. In other words, regions with slightly weaker match between the RFMix and FLARE inferences cannot be resolved to be more accurate in one compared to the other.

## Materials and Methods

### The HCHS/SOL

The Hispanic Community Health Study / Study of Latinos (HCHS/SOL) is a multi-center prospective study (2008-2011) for individuals of Hispanic/Latino origin aged 18-74 years old at recruitment. The study aims to identify the prevalence and factors impacting specific chronic conditions such as cardiovascular diseases, asthma, and sleep disorders in Hispanic/Latino individuals [48]. Participants were recruited from four US field centers: Bronx (New York), Chicago (Illinois), San Diego (California), and Miami (Florida) [5,48]. Individuals were sampled for the study via a two-stage probability sample of household in census-block groups, which were predefined by communities that are enriched for Hispanics/Latinos near the study centers. Each study participant self-reported their Hispanic/Latino background, with major groups being Cuba, Dominican Republic, Mexico, Puerto Rico, Central America, and South America [30,31].

Demographic information (sex and age at the time of participant’s clinic visit), clinical measures including body mass index (BMI), and the estimated glomerular filtration rate (eGFR) were recorded at baseline [9]. EGFR levels were estimated based on serum cystatin C without demographic factors using the equation developed from Chronic Kidney Disease Epidemiology Collaboration [49].

### Genotyping, imputation, and relatedness inference

HCHS/SOL individuals from the previously-reported RFMix-inferred local ancestry dataset were genotyped using the Illumina Omni2.5M SNVs array, which included ∼150K custom SNPs designed to capture genetic variation relevant to Hispanic/Latino individuals [27]. The RFMix-inferred dataset uses genome build GRCh37 (hg19) coordinates. Details about genotyping, phasing, imputation, and relatedness inference for this dataset has been published [35]. The HCHS/SOL individuals in the FLARE newly-inferred datasets (described below) were genotyped using the Multi-Ethnic Genotyping Array (MEGA), as part of the HCHS/SOL participations in the Population Architecture Using Genomics and Epidemiology (PAGE) consortium [50]. The genotyped data were then imputed using The Trans-Omics in Precision Medicine (TOPMed) 1.0 reference panel [51]. Both reference haplotypes and HCHS/SOL variant data were phased using Beagle v.5.4 before applying FLARE [52]. In the FLARE files, SNPs were retained for analysis if their minor allele frequency (MAF)≥ 0.005 and (if imputed) imputation quality score R^2^≥ 0.8 as a high and run-time efficient imputation quality threshold [20,53]. The FLARE-inferred datasets use genome build GRCh38 (hg38) coordinates [20,53].

All association analyses used principal components (PCs), and kinship matrix computed based on the Omni2.5M array genotyping [35]. Genetic PCs and kinship matrix were computed using the KING-robust [54] for initial kinship coefficient estimates, and then PC-AiR [55], and PC-relate[56] algorithms implemented in the GENESIS R package [57] as previously reported [35]. This prior work demonstrated the first 5 PCs are sufficient to account for population structure in association analysis for HCHS/SOL and developed “genetic analysis groups”, groups of individuals that are largely overlapping their self-reported Hispanic/Latino background but modified so that each group is genetically homogeneous on the Euclidian space defined by the first 5 PCs. These groups assigned individuals with “other” or missing Hispanic/Latino background into one of the 6 major groups (Central American, Cuban, Dominican, Mexican, Puerto Rican, and South American). The term “genetic analysis group” reflects the use of these grouping solely for genetic analysis purposes and should not be confused with self-reported Hispanic/Latino ethnic background.

### Local ancestry inference: RFMix and FLARE

RFMix local ancestry was inferred using reference genotypes from three continents (Africa, America, and Europe). The reference panel was constructed using data from the Human Genome Diversity Project (HGDP) and the 1000 Genomes Project (1000G), comprising 195 West African, 63 Amerindian, and 527 European participants genotyped with either the Illumina HumanHap650Y array (HGDP) or the Omni2.5M array (1000G) [19,34]. Participants who had a whole-chromosome anomaly on any of the autosomes were excluded from the autosomal local ancestry calculation. The detailed process of genotyping, imputations, and local ancestry calls, were published [5,27]. The inferred dataset includes 12,689 individuals and local ancestry counts over 15,500 intervals. Local ancestry inferences were conducted on both the autosomes and the X chromosome; however, only the autosomes were included in the analysis to align with the FLARE dataset.

FLARE-inferred datasets include FLARE7 (using 7 reference populations) and FLARE3 (using 3 reference populations). Both local ancestry inferences used FLARE v0.3.0 and reference haplotypes generated from the Human Genome Diversity Project (HGDP) data [34], which were derived from whole-genome sequencing data of 929 participants across seven geographic regions: Sub-Saharan Africa (104), Central-South America (61), Europe (155), Central/South Asia (197), East Asia (223), Oceania (28), and the Middle East (161). The FLARE7 dataset comprises 11,928 individuals and 5,105,005 SNPs, utilizing reference populations from all seven regions. In contrast, FLARE3 inference was based on reference populations from three regions (Africa, America, and Europe) and includes the same 11,928 HCHS/SOL individuals but 3,618,751 SNPs. The different number of SNPs is due to the difference in reference populations used. The FLARE local ancestry inference includes a smaller set of HCHS/SOL individuals compared to the RFMix inferred local ancestries because it used the subset of HCHS/SOL individuals genotyped via the MEGA array. Subsequent comparisons between three inferences used individuals participating in all datasets (n=11,863). Unlike the RFMix-based local ancestry calls, the FLARE7 and FLARE3 datasets provided local ancestry calls at the SNP level, without merging into intervals.

### Global ancestry proportions: RFMix, FLARE, and ADMIXTURE

We computed global ancestry proportions based on the inferred local ancestry counts. For a given individual, the RFMix ancestry inference, the global proportion for each ancestry was calculated by summing the lengths of all local ancestry intervals, multiplying each by the corresponding ancestry counts, and then dividing the resulting number by twice the total length of all intervals (since the local ancestry counts can be at most 2). For files derived from the FLARE analysis, the global proportion for each ancestry was calculated by taking the equally weighted average of local ancestry counts across all SNPs: computing half of the mean local ancestry counts for each participant.

We also compared the above global ancestry proportions to those previously computed using a supervised ADMIXTURE [36] model over a set of unrelated HCHS/SOL individuals, under the assumption of 3 ancestral populations (African, Amerindian, and European, k = 3). The details of reference population generation, relatedness thresholding, and SNPs selection were published[35].

### Comparison of inferred local ancestries

The previous, RFMix, inference was conducted based on Human Genome Assembly GRCh37 (hg19), while the new inferences FLARE3 and FLARE7 were based on GRCh38 (hg38). We used the LiftOver tool on UCSC Genome Browser [38,58] to align the RFMix-inferred intervals to hg38. Intervals that were mapped into more than one region or that were mapped to a different chromosome were discarded.

SNPs in the FLARE3 and FLARE7 inferences were matched to the local ancestry intervals from the RFMix inference separately. A SNP was matched to an interval in the RFMix if the SNP position was greater or equal to the block starting position and less or equal to the block ending position. For each variant-interval pair, and for each of European, African, and Amerindian ancestries, Pearson correlation was computed to assess the consistency of the local ancestry counts among the inferences across the genome (chromosome 1-22) across all individuals.

Regions with relatively low local ancestry correlation between the RFMix and the FLARE datasets (< 0.9) were aligned to annotations from UCSC genome browser, which provides coordinates of well-known gene clusters that can yield alignments with low-quality mapping scores and discordant read pairs [38], and to the ENCODE “blacklist”, which comprises areas exhibiting anomalous, unstructured, or high signal in next-generation sequencing experiments, regardless of experiments [59]. To assess whether associations of low correlations between RFMix and FLARE local ancestries are enriched in these annotated problematic regions, we also selected at random a matching number of regions with high correlations (>0.97) between RFMix and FLARE local ancestries and compared the number of regions aligned to the annotated problematic regions between the low and high correlation groups. Even though genotyping arrays were used, the reference panels and reference haplotypes used in imputation and local ancestry inference incorporated whole genome and whole exome sequencing data [19,51,60], therefore could potentially propagate the bias of the blacklisted regions into the local ancestry calls. Because local ancestry has long range linkage disequilibrium (LD), we allowed up to 0.5 Mb threshold to determine that a region with low correlation in inferred local ancestry counts is associated with a blacklisted interval.

### Metabolomics data

Metabolomics data were assayed in fasting serum in two separate profiling efforts in HCHS/SOL. First, in 2017, 4,002 randomly-selected HCHS/SOL participants who also underwent genotyping were assayed (we refer to this dataset as “batch 1”). Next, in 2021, additional 2,368 serum samples from 2,330 participants were assayed using blood samples collected at baseline (“batch 2”) [27]. Batch 2 included repeated samples within it, as well as samples from individuals who were assayed in batch 1, for quality control. Batch 2 participants also included individuals who were sampled for various ancillary studies: it included (1) individuals who participated in the ECHO-SOL ancillary study of echocardiogram [61], (2) individuals who had normal estimated glomerular filtration rate (eGFR > 60 at the baseline HCHS/SOL exam and substantial decline from the baseline to the second HCHS/SOL exam, and (3) individuals with eGFR measures available from both the baseline and second HCHS/SOL exams.

Serum samples were stored at −70°C at the HCHS/SOL Core Laboratory at the University of Minnesota until analysis by Metabolon, Inc. (Durham, NC) in 2017 (batch 1) and 2021 (batch 2). Serum samples were then extracted and prepared using Metabolon’s standard solvent extraction method. Extracts were split into five fractions to use in four liquid chromatography-mass spectrometry (LC-MS)-based metabolomic quantification platforms (two reverse phase methods with positive ion mode electrospray ionization (EI), one reverse phase method with negative ion mode EI, and one hydrophilic interaction liquid chromatography with negative ion mode EI), with the fifth fraction reserved for backup. Instrument variability was assessed by calculating the median relative standard deviation (SD) for the internal standards added to each sample prior to injection into the mass spectrometers. Overall process variability was determined by calculating the median relative SD for all endogenous metabolites (i.e., non-instrument standards) present in 100% of the technical replicate samples.

### Comparison of admixture mapping of metabolites in previously-reported association regions

In previous work, the two non-overlapping datasets (discovery and replication datasets) extracted from metabolomics batch 1 and batch 2 were used to examine genetic associations with 640 metabolites via admixture mapping [27]. Four of the metabolites with the strongest admixture mapping associations reported (according to the association p-value) were: 3-aminoisobutyrate, N-acetylarginine, PE 16:0/20:4, and PC 16:0/20:4 [27]. For both metabolomics datasets, individuals with complete data for the four metabolites and who also are available in the FLARE and RFMix datasets were included in admixture mapping analyses (batch 1 discovery: n = 3861, batch 2 replication: n = 1644).

Admixture mapping between each considered metabolite and ancestry counts across the genome was tested with Score tests using R package GENESIS [57]. We focused on African and Amerindian ancestries, where each was tested separately using local ancestry counts from RFMix, FLARE3, and FLARE7 inferences. Null models were fitted first, i.e. models without genetic ancestry count, adjusting for fixed covariate effects (age, sex, center, eGFR, genetic analysis group, and the first five genetic PCs) as well as random effects accounting for the study design of HCHS/SOL and relatedness (household, census-block unit, and kinship coefficients) [27]. Genome-wide multiple testing burden was computed using STEAM R package [62], with significance thresholds determined for each inference and each batch based on the genetic distances of a sampled subset of pairs of SNPs obtained from the HapMap GRCh38 genetic map [63], correlations between local ancestry counts for each pair of sampled SNPs, and individual global ancestry proportions computed from local ancestry counts as described earlier. When constructing SNPs correlation matrices using the FLARE7 dataset, the correlations among the 7 ancestries were simplified to the 3 major ancestries (African, Amerindian and European) and one combined local ancestry counts for the other four ancestries. This simplification was conducted due to the high sparsity of some ancestries, which made their individual local ancestry counts less informative for SNPs correlation and the downstream inference of the number of generations since admixture.

To assess replicability of the analyses between the discovery and replication batches in each local ancestry inference, we first identified the most significantly associated genomic regions using RFMix inference for each metabolite and ancestry in the discovery dataset (taken from batch 1). Then within each specific genomic region, the p-values and rsIDs of the most significantly associated variant using FLARE3 and FLARE7 were identified, and the association effect sizes as well as p-values were compared across different local ancestry inferences and the two batches.

## Acknowledgements

The authors thank the staff and participants of HCHS/SOL for their important contributions to the HCHS/SOL studies. A complete list of staff and investigators has been provided by Sorlie P., et al. Investigators in Ann Epidemiol. 2010 Aug;20: 642-649 and is also available on the study website - http://www.cscc.unc.edu/hchs/. The Hispanic Community Health Study/Study of Latinos is a collaborative study supported by contracts from the National Heart, Lung, and Blood Institute (NHLBI) to the University of North Carolina (HHSN268201300001I / N01-HC-65233), University of Miami (HHSN268201300004I / N01-HC-65234), Albert Einstein College of Medicine (HHSN268201300002I / N01-HC-65235), University of Illinois at Chicago (HHSN268201300003I / N01-HC-65236 Northwestern Univ), and San Diego State University (HHSN268201300005I / N01-HC-65237). The following Institutes/Centers/Offices have contributed to the HCHS/SOL through a transfer of funds to the NHLBI: National Institute on Minority Health and Health Disparities, National Institute on Deafness and Other Communication Disorders, National Institute of Dental and Craniofacial Research, National Institute of Diabetes and Digestive and Kidney Diseases, National Institute of Neurological Disorders and Stroke, NIH Institution-Office of Dietary Supplements. This study was supported by National Human Genome Research Institute grants R56HG013163 and R01HG013163.

## Ethics statement

The HCHS/SOL was approved by the institutional review boards (IRBs) at each field center, where all participants gave written informed consent, and by the Non-Biomedical IRB at the University of North Carolina at Chapel Hill, to the HCHS/SOL Data Coordinating Center. All IRBs approving the HCHS/SOL study are: Non-Biomedical IRB at the University of North Carolina at Chapel Hill. Chapel Hill, NC; Einstein IRB at the Albert Einstein College of Medicine of Yeshiva University. Bronx, NY; IRB at Office for the Protection of Research Subjects (OPRS), University of Illinois at Chicago. Chicago, IL; Human Subject Research Office, University of Miami. Miami, FL; Institutional Review Board of San Diego State University, San Diego, CA. All methods and analyses of HCHS/ SOL participants’ materials and data were carried out in accordance with human subject research guidelines and regulations. This work was approved by the Beth Israel Deaconess Medical Center Committee on Clinical Investigations.

## Data and code availability statement

HCHS/SOL data are available through application to the data base of genotypes and phenotypes (dbGaP) accession phs000810, or via a data use agreement with the HCHS/SOL Data Coordinating Center (DCC) at the University of North Carolina at Chapel Hill, see collaborators website: https://sites.cscc.unc.edu/hchs/. The ENCODE blacklist (hg38) can be accessed through the GitHub repository at https://github.com/Boyle-Lab/Blacklist/tree/master/lists, while the UCSC unusual regions data, which includes gene cluster information, is available for download at https://hgdownload.soe.ucsc.edu/gbdb/hg38/problematic/comments.bb. Code used for analysis in this paper is publicly available on the GitHub repository https://github.com/xc448/HCHS-SOL_local_ancestry_comparison

## Author contribution statement

TS conceptualized the study. HW performed FLARE analysis. XC performed all comparisons between global and local ancestries, admixture mapping, prepared tables and figures. XC drafted the manuscript. All authors critically reviewed the manuscript. TS supervised the work.

