## Supplementary Materials for "Old vs. New Local Ancestry Inference in HCHS/SOL: A Comparative Study"

Supplementary Information

### Supplementary Figures

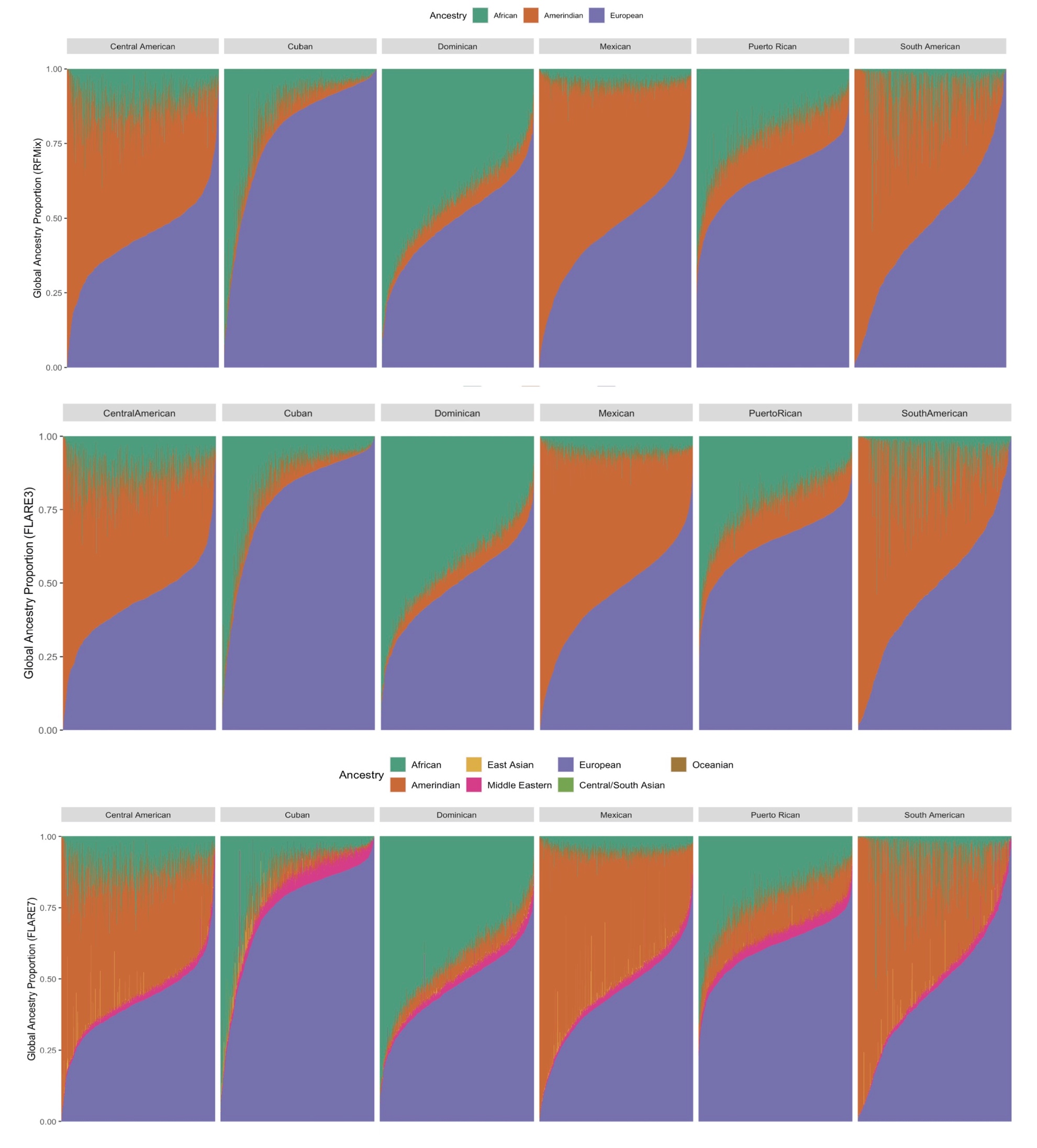

#### Supplementary Figure 1: Global Ancestry Proportions Computed from RFMix, FLARE3, and FLARE7 Dataset

**(B)**

**(C)**

**(A)**

Global ancestry proportions estimated by RFMix (A), FLARE3 (B), and FLARE7 (C), sorted by proportions of European ancestry and visualized by previously defined “genetic analysis groups”.

#### Supplementary Figure 2: Values of the first two genetic PCs of HCHS/SOL participants colored by global ancestry proportions

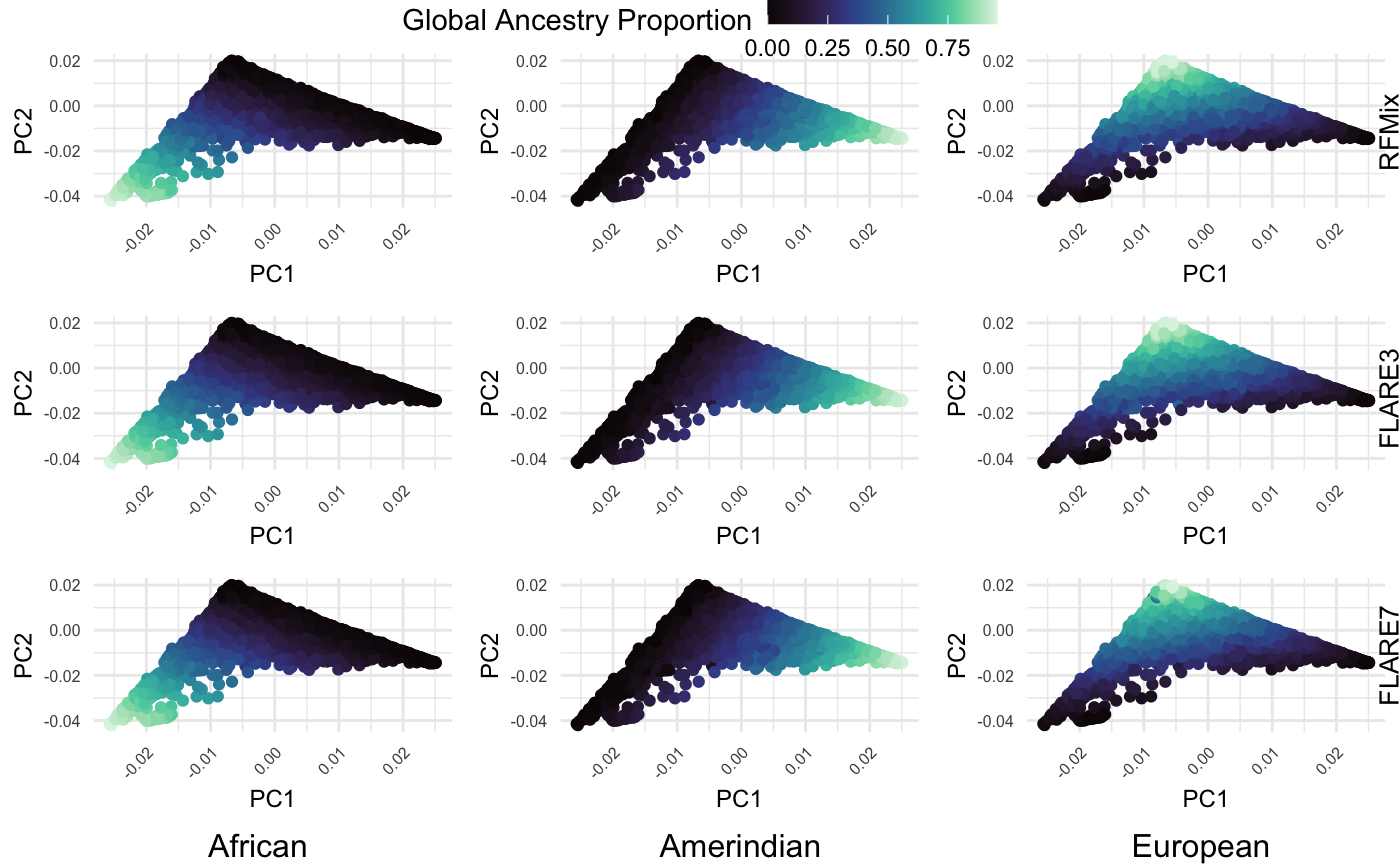

The first two genetic PCs plotted against each other and colored by global ancestry proportions (African, Amerindian, and European ancestries) estimated from RFMix (top), FLARE3 (middle), and FLARE7 (bottom).

#### Supplementary Figure 3: Pearson correlation of local ancestry counts with SNPs filtered by R^2^ $\geq$ 0.95

**(B)**

**(A)**

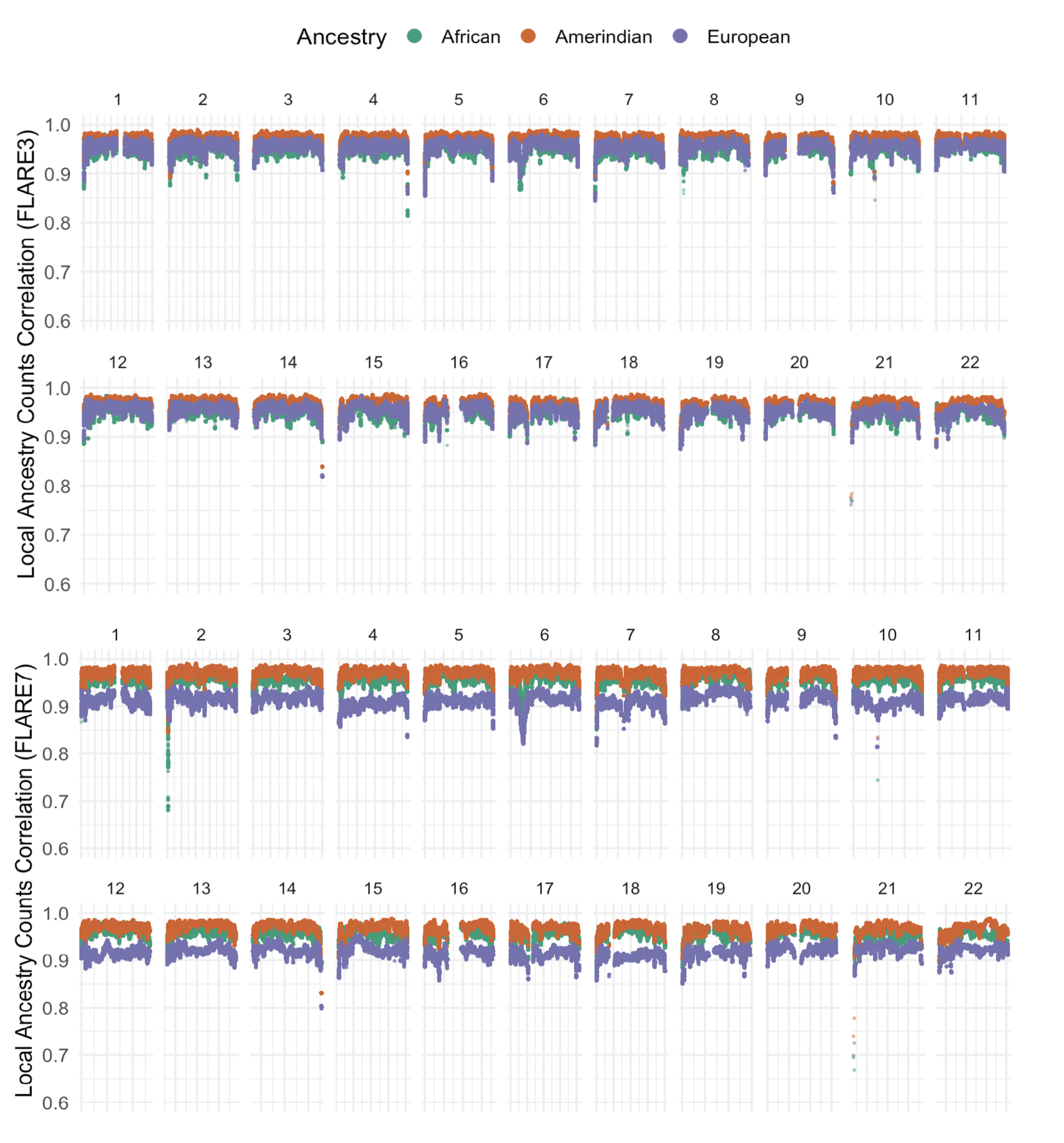

Pearson correlation of the local ancestry counts between FLARE3 and RFMix (A) and FLARE7 and RFMix (B) for each matched SNP and ancestry block across chromosome 1-22 with all SNPs with imputation quality threshold R^2^ $\geq$0.95.

#### Supplementary Figure 4: Mean ancestry proportions across HCHS/SOL individuals for each SNP (FLARE3) or interval (RFMix).

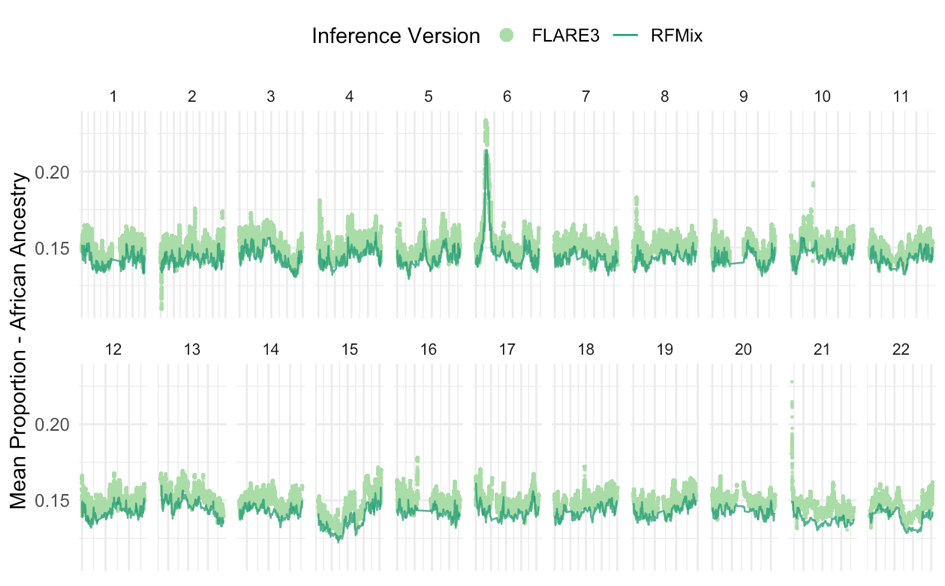

**(A)**

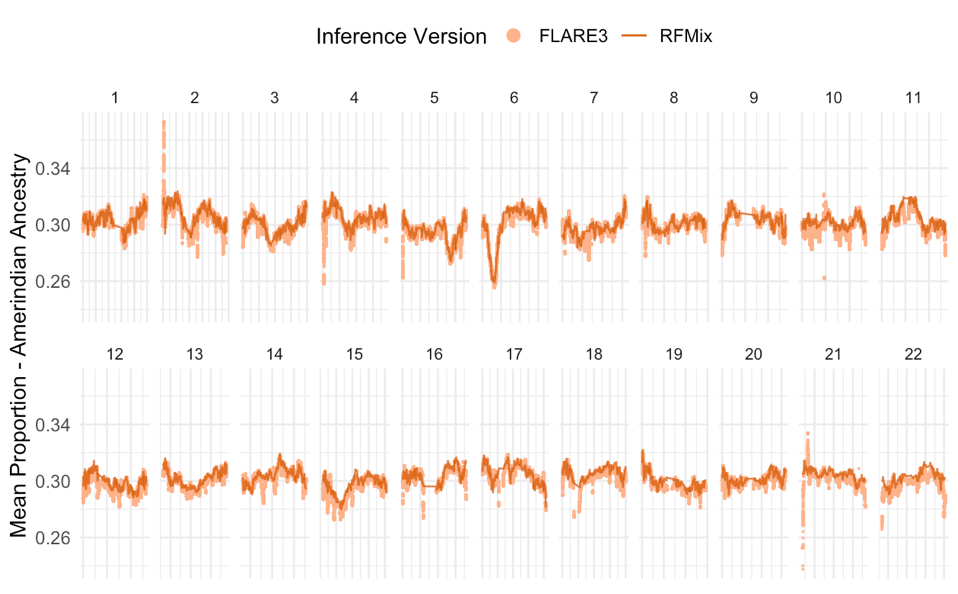

**(B)**

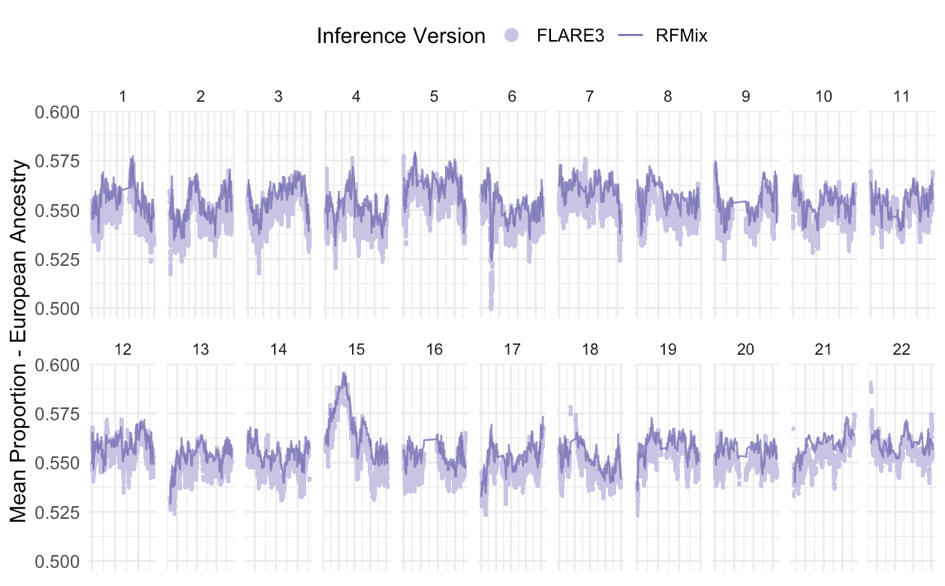

**(C)**

Mean ancestry proportions (African, Amerindian, and European ancestry) according to RFMix and FLARE3 inference for each matched SNP and ancestry block across chromosome 1-22 across all HCHS/SOL individuals with genetic data.

#### Supplementary Figure 5: Mean ancestry proportions across HCHS/SOL individuals for each SNP (FLARE7) or ancestry interval (RFMix)

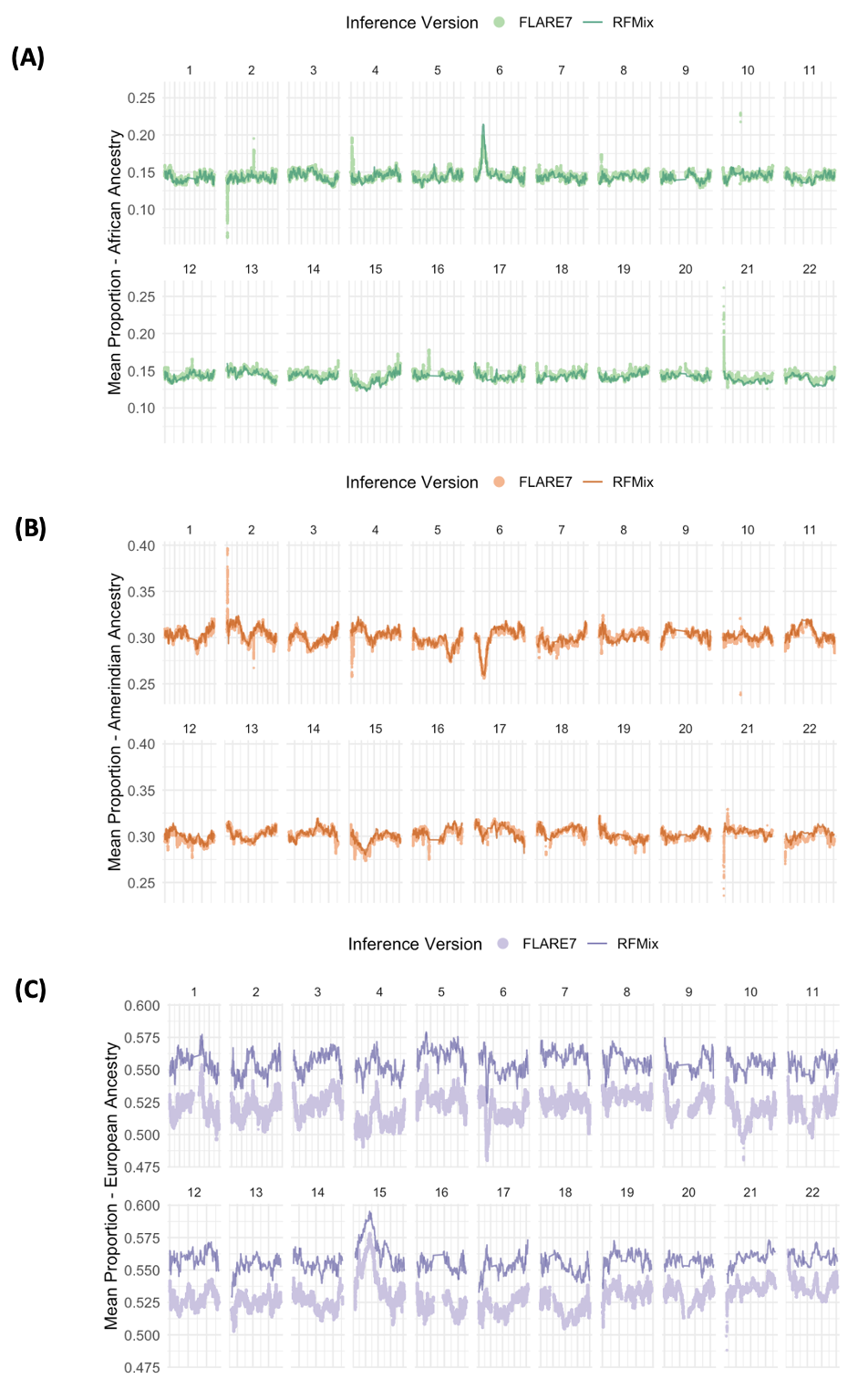

Mean ancestry proportions (African, Amerindian, and European ancestry) according to RFMix and FLARE7 inference for each matched SNP and ancestry interval across chromosome 1-22 across all HCHS/SOL individuals.

#### Supplementary Figure 6: Variance of ancestry proportions across HCHS/SOL individuals for each SNP (FLARE3) or ancestry interval (RFMix)

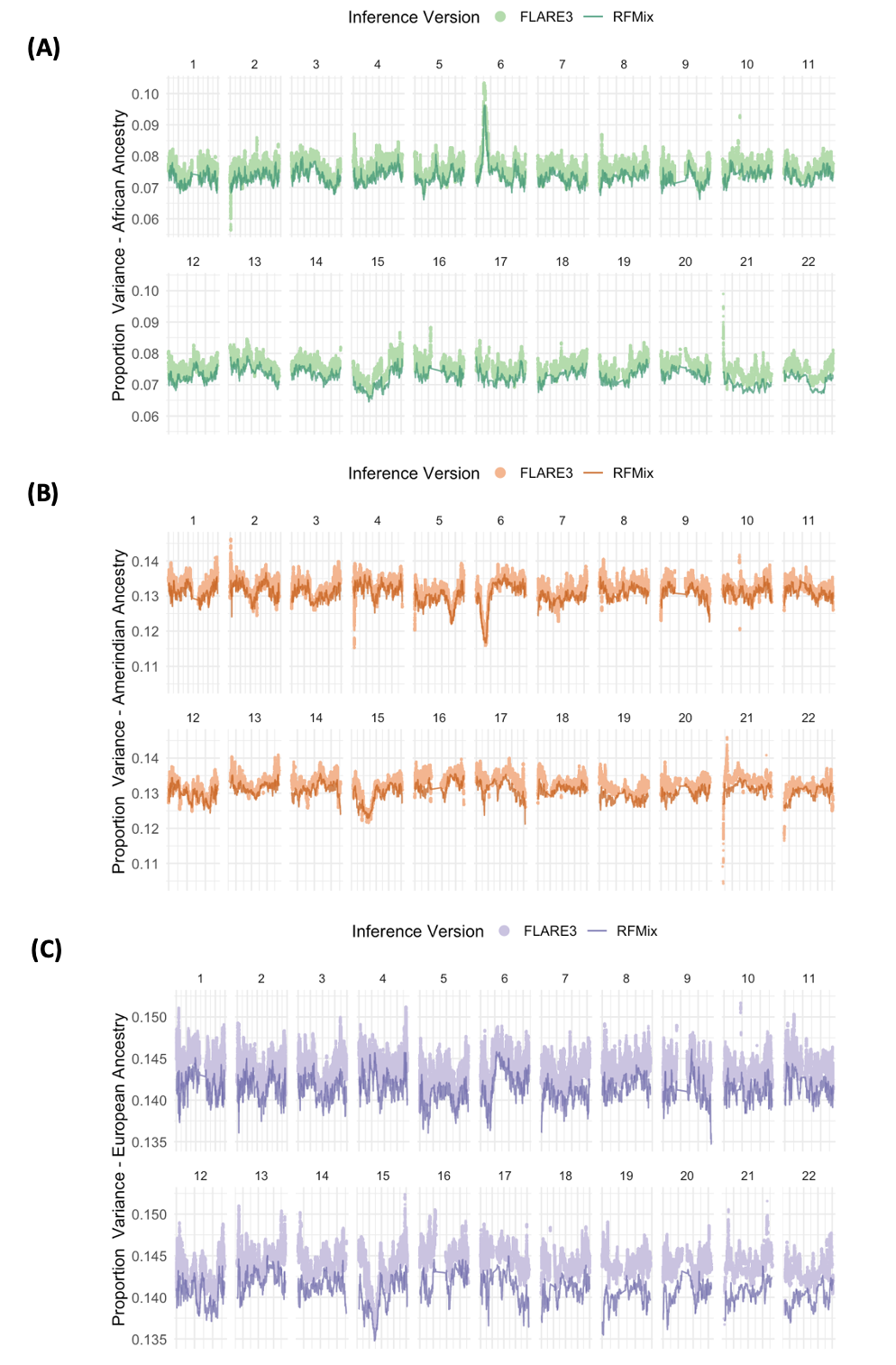

Variance of ancestry proportions (African, Amerindian, and European ancestry) according to the RFMix and FLARE3 inference for each matched SNP and ancestry block across chromosome 1-22 across all HCHS/SOL individuals.

#### Supplementary Figure 7: Variance of ancestry proportions across HCHS/SOL individuals for each SNP (FLARE7) or ancestry interval (RFMix)

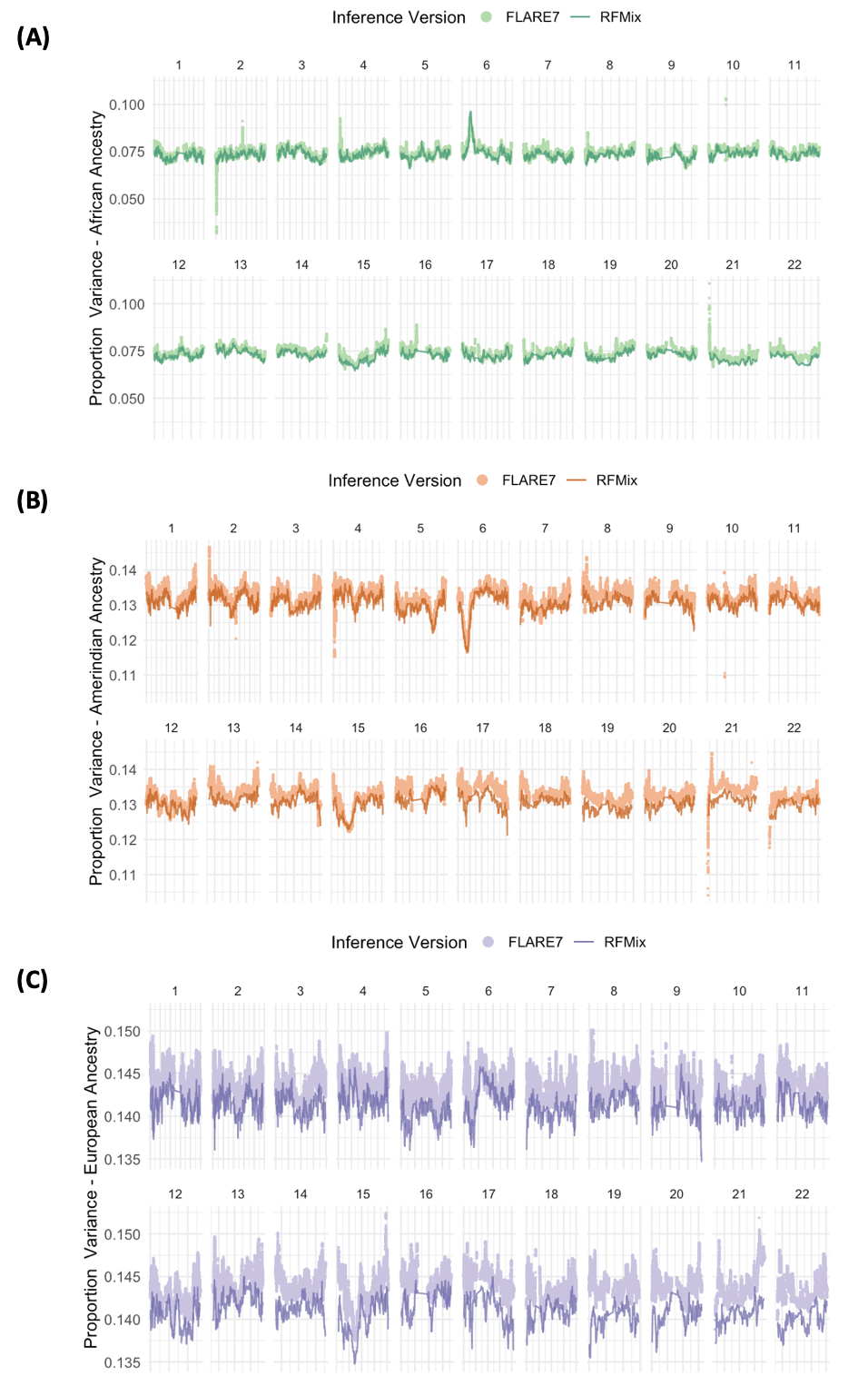

Variance of ancestry proportions (African, Amerindian, and European ancestry) according to RFMix and FLARE7 inference for each matched SNP and ancestry block across chromosome 1-22 across all HCHS/SOL individuals.

#### Supplementary Figure 8: Regional association plot for admixture mapping of N-acetylarginine, focusing on African ancestry at chromosome 2

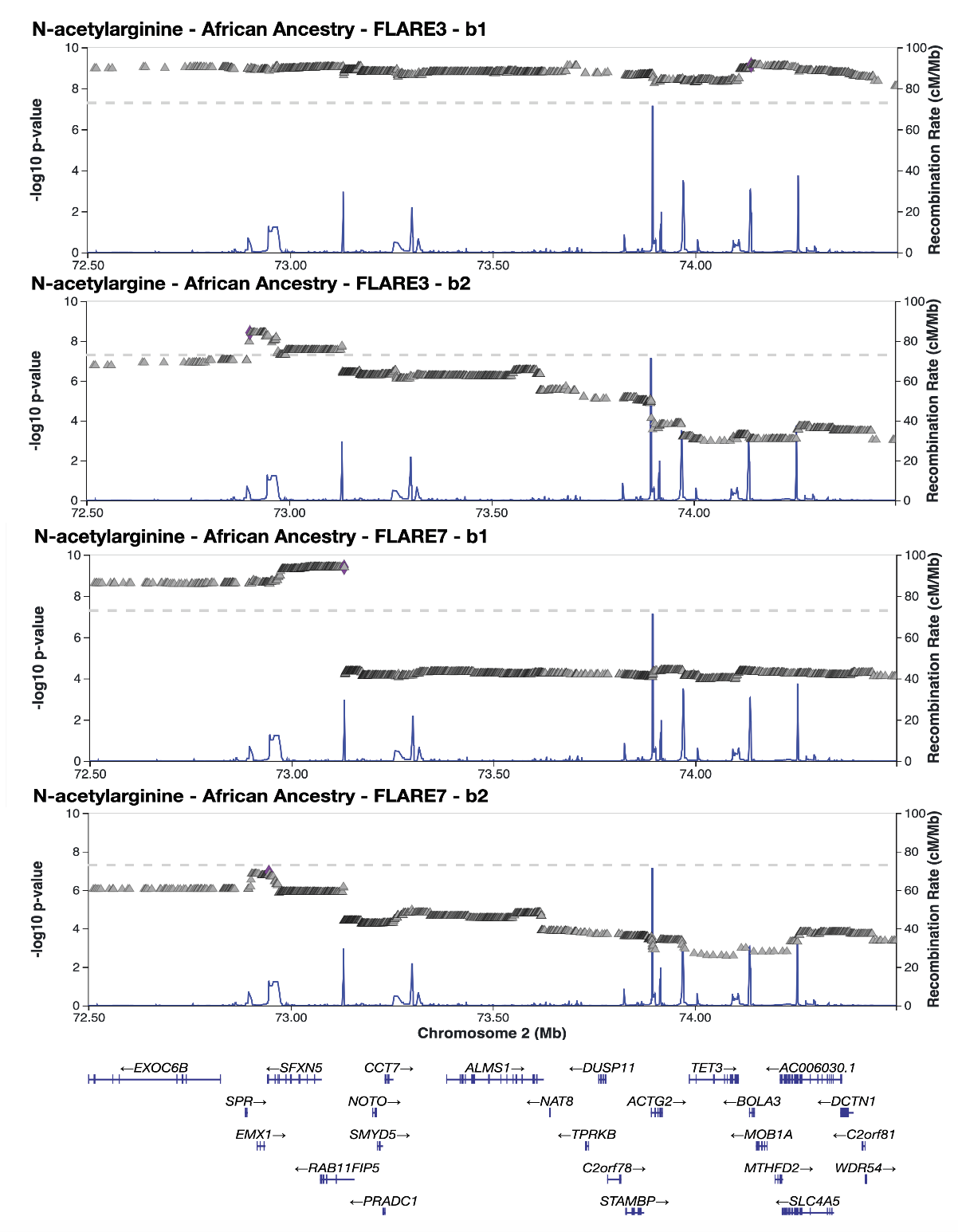

**(C)**

**(D)**

**(B)**

**(A)**

Zoomed-in view of SNPs with -log10 p-values (inverted triangles) in the top region of chromosome 2 from admixture mapping of N-acetylarginine using African local ancestry from FLARE3 (A, B) or FLARE7 (C, D) inference across batches. Blue lines indicate recombination rates, centered on the reported gene ALMS1, with arrows highlighting the most significant SNP (purple diamonds).

#### Supplementary Figure 9: Regional association plot for admixture mapping of 3-aminoisobutyrate focusing on Amerindian ancestry at chromosome 5

**(A)**

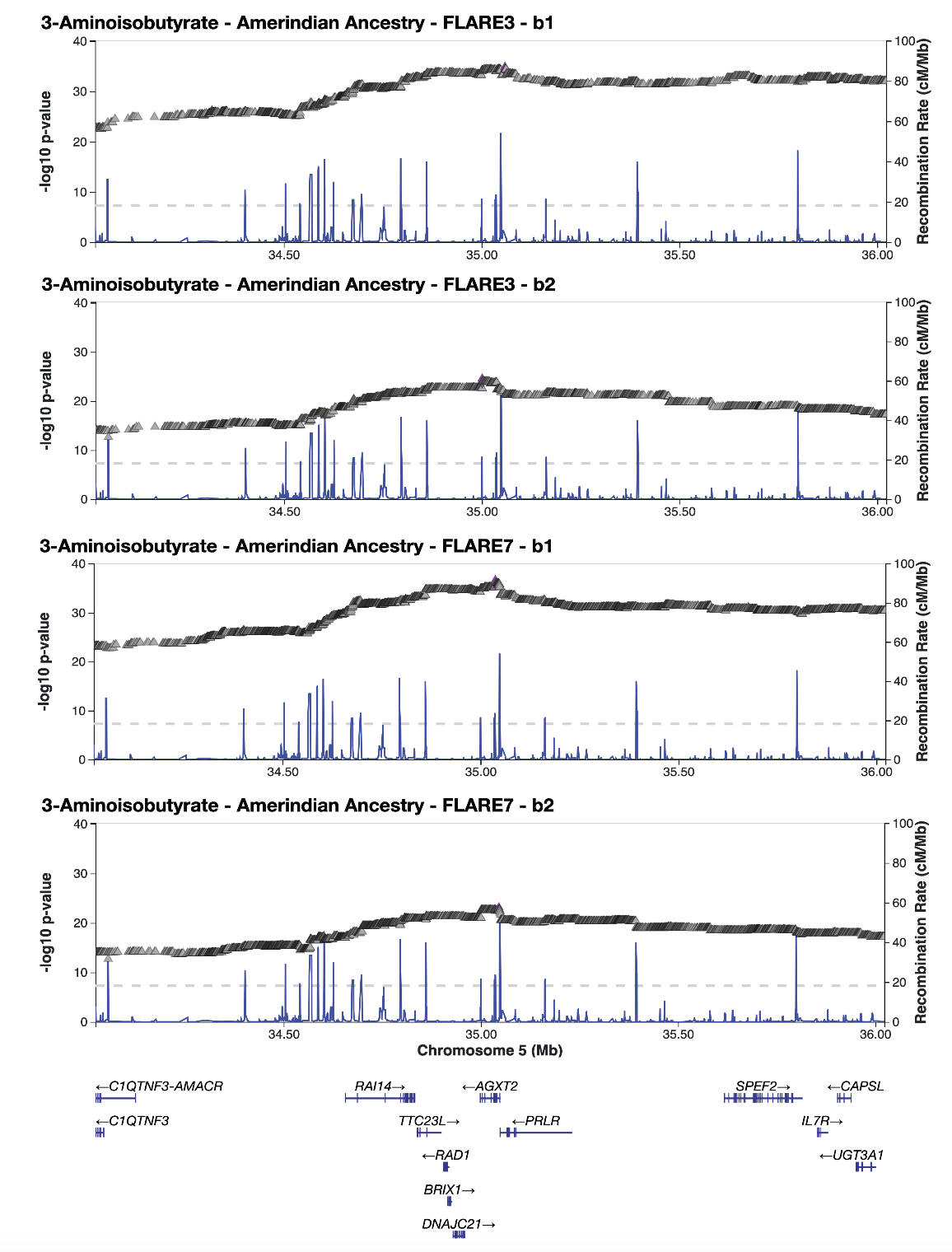

**(D)**

**(C)**

**(B)**

Zoomed-in view of SNPs with -log10 p-values (inverted triangles) in the top region of chromosome 5 from admixture mapping of 3-aminoisobutyrate using Amerindian local ancestry from FLARE3 (A, B) or FLARE7 (C, D) inference across batches. Blue lines indicate recombination rates, centered on the reported gene AGXT2, with arrows highlighting the most significant SNP (purple diamonds).

#### Supplementary Figure 10: Regional association plot for admixture mapping of PC 16:0/20:4, focusing on African ancestry at chromosome 11

**(C)**

**(A)**

**(D)**

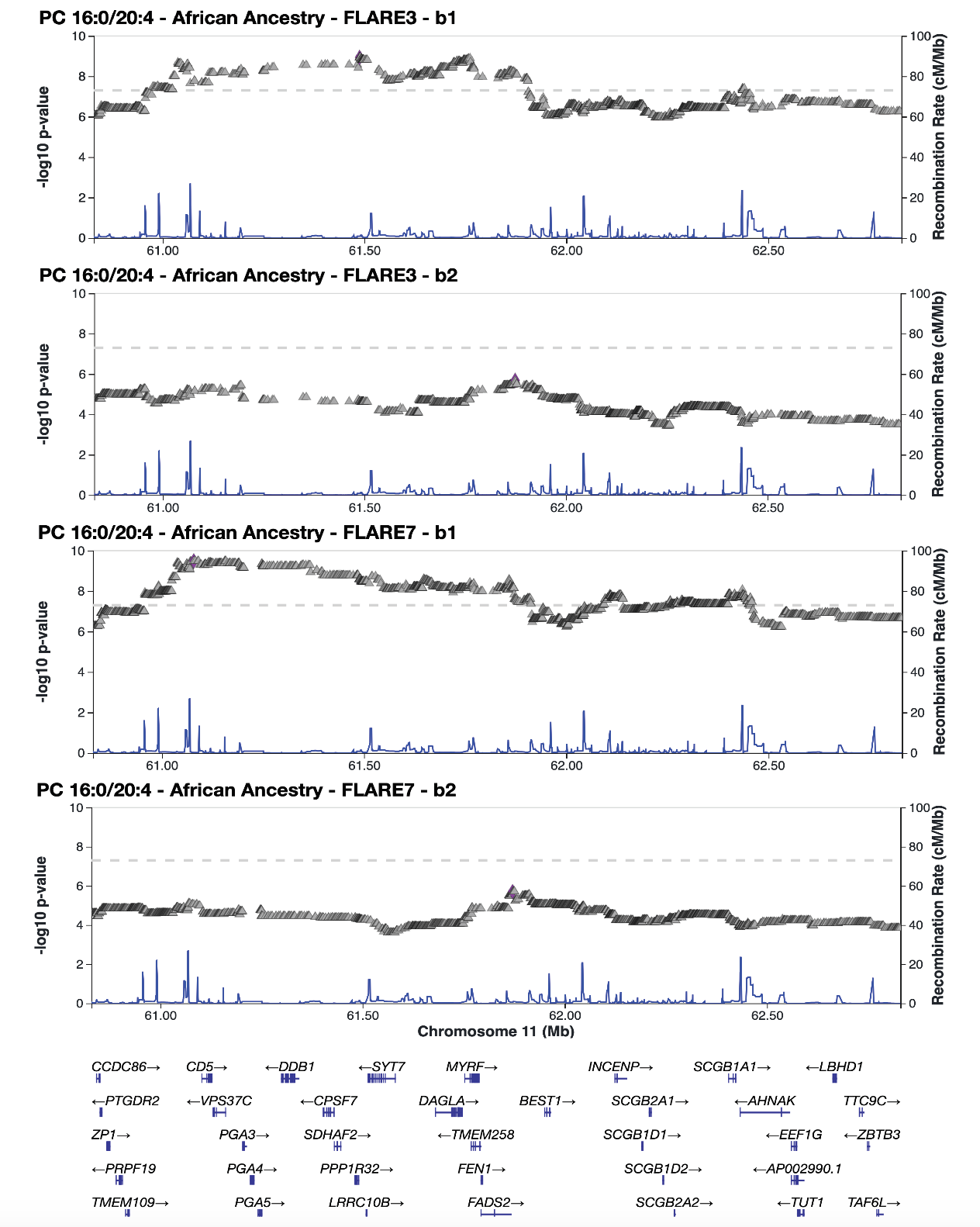

**(B)**

Zoomed-in view of SNPs with -log10 p-values (inverted triangles) in the top region of chromosome 11 from admixture mapping of PC 16:0/20:4 using African local ancestry from FLARE3 (A, B) or FLARE7 (C, D) inference across batches. Blue lines indicate recombination rates, centered on the reported gene FADS2, with arrows highlighting the most significant SNP (purple diamonds).

#### Supplementary Figure 11: Regional association plot for admixture mapping of PC 16:0/20:4, focusing on Amerindian ancestry at chromosome 11

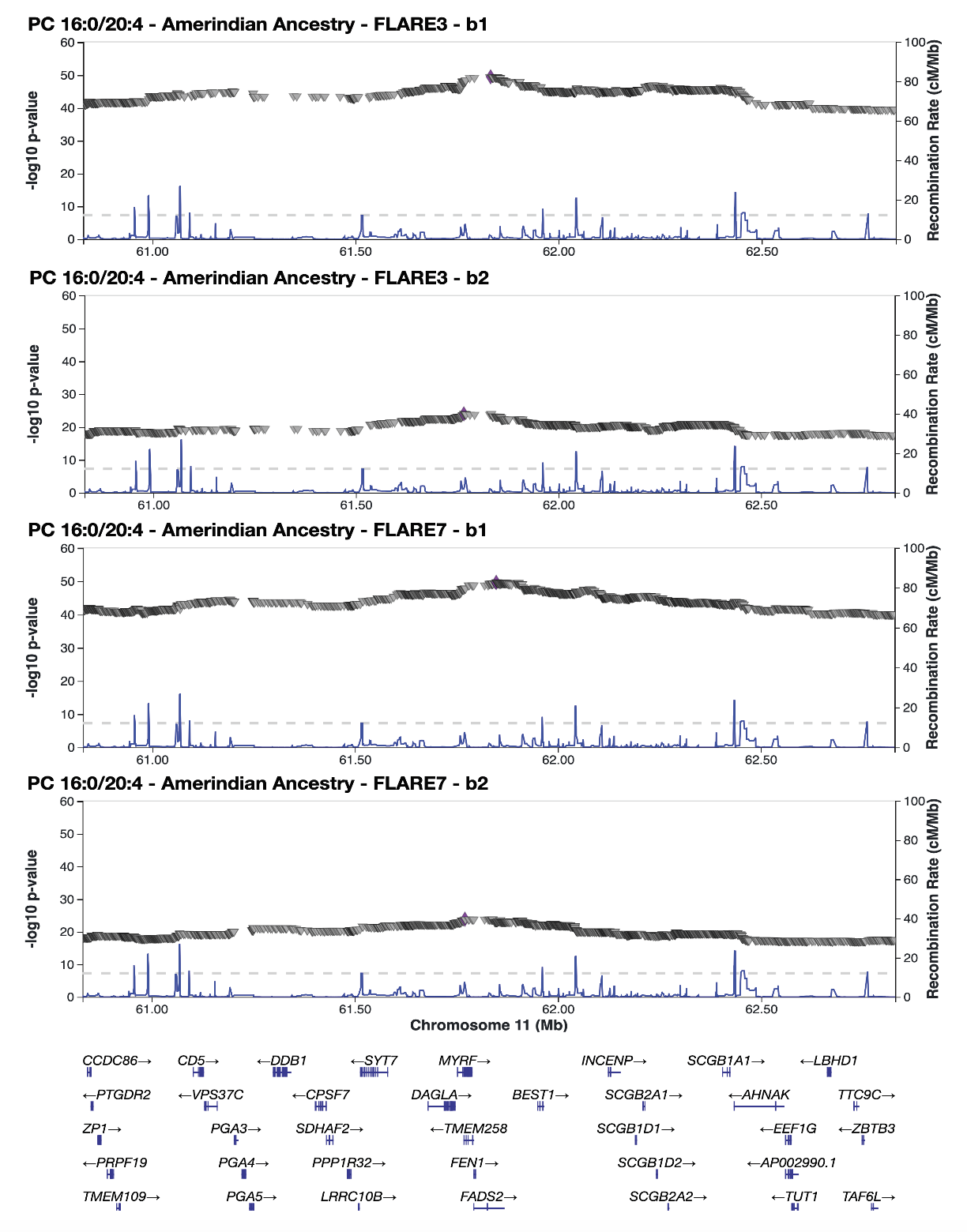

**(D)**

**(C)**

**(B)**

**(A)**

Zoomed-in view of SNPs with -log10 p-values (inverted triangles) in the top region of chromosome 11 from admixture mapping of PC 16:0/20:4 using American local ancestry from FLARE3 (A, B) or FLARE7 (C, D) inference across batches. Blue lines indicate recombination rates, centered on FADS2, with arrows highlighting the most significant SNP (purple diamonds).

#### Supplementary Figure 12: Regional association plot for admixture mapping of PE 16:0/20:4, focusing on Amerindian ancestry at chromosome 15

**(B)**

**(D)**

**(C)**

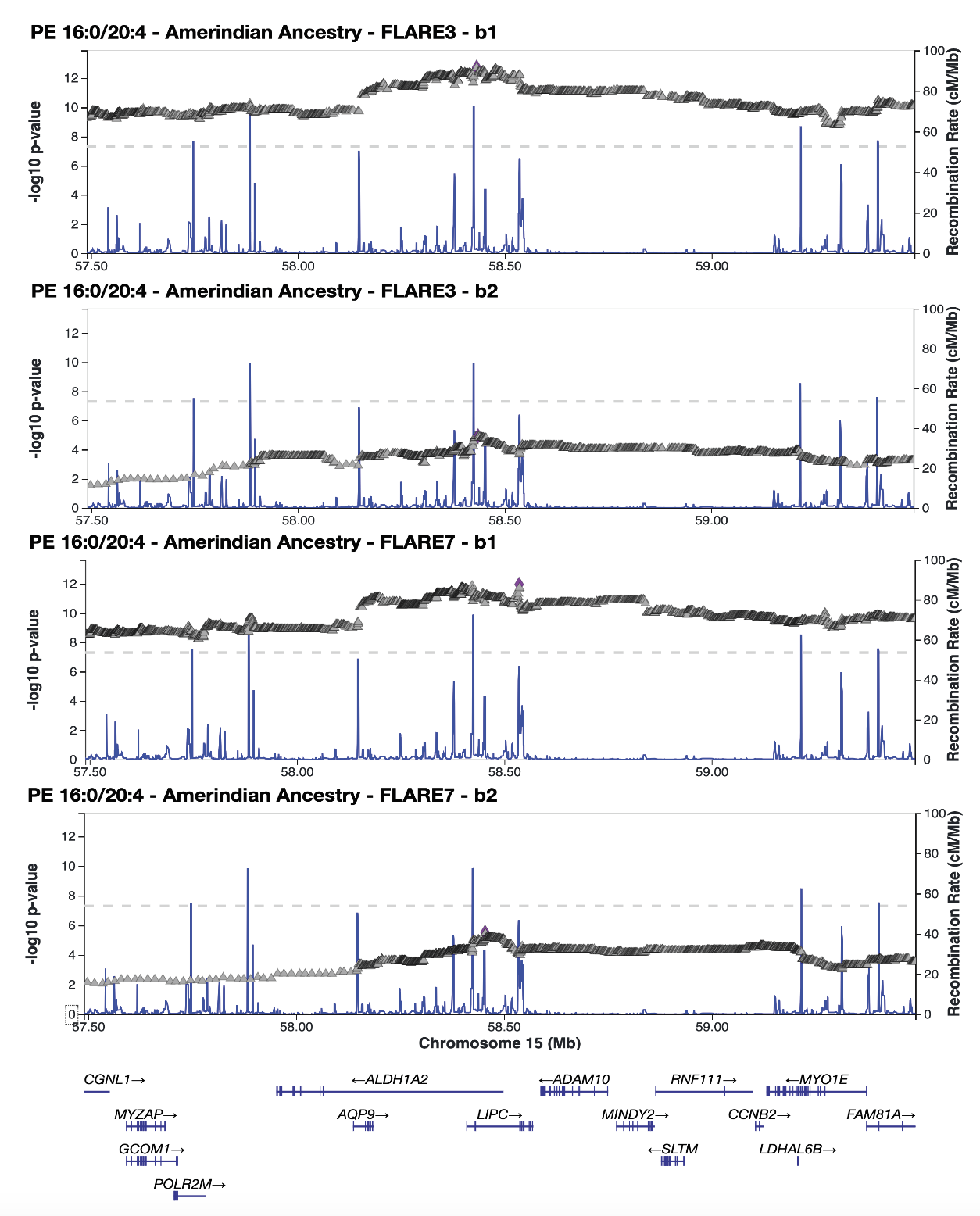

**(A)**

Zoomed-in view of SNPs with -log10 p-values (inverted triangles) in the top region of chromosome 15 from admixture mapping of PE 16:0/20:4 using American local ancestry from FLARE3 (A, B) or FLARE7 (C, D) inference across batches. Blue lines indicate recombination rates, centered on LIPC, with arrows highlighting the most significant SNP (purple diamonds).

#### Supplementary Figure 13: Admixture mapping result of propyl 4-hydroxybenzoate sulfate, focusing on African ancestry at chromosome 16

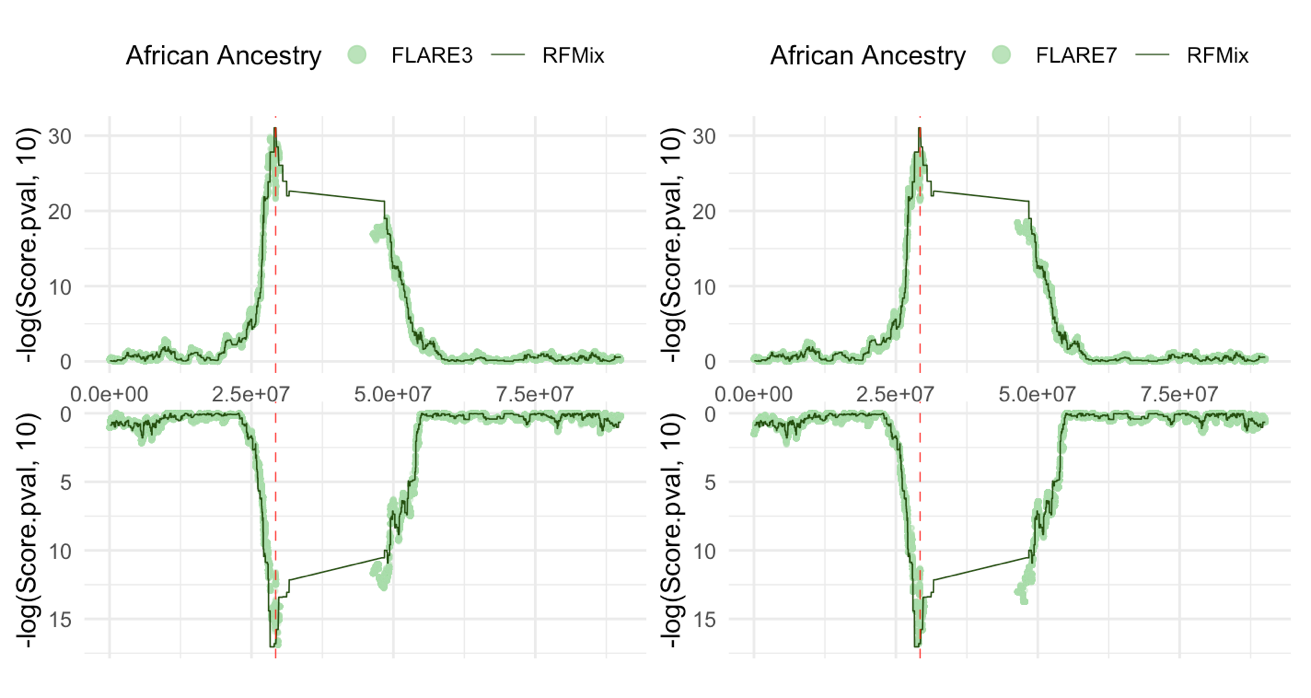

Admixture mapping using local ancestry counts in RFMix vs. FLARE3 (left) and RFMix vs. FLARE7 (right) at the most significant chromosome for the metabolite propyl 4-hydroxybenzoate sulfate at chromosome 16. The results from the two batches are displayed as mirrored plots, with the upper panel representing batch 1 and the lower panel representing batch 2. The red dashed lines indicate the location of low correlation (between local ancestry counts in RFMix and FLARE inferences) region mapped to the ENCODE blacklist.

### Supplementary Tables

#### Supplementary Table 1: Aligned regions of ENCODE blacklist or annotations of gene clusters from UCSC genome browser to the low correlation SNP-LAI pair between FLARE3 and RFMix

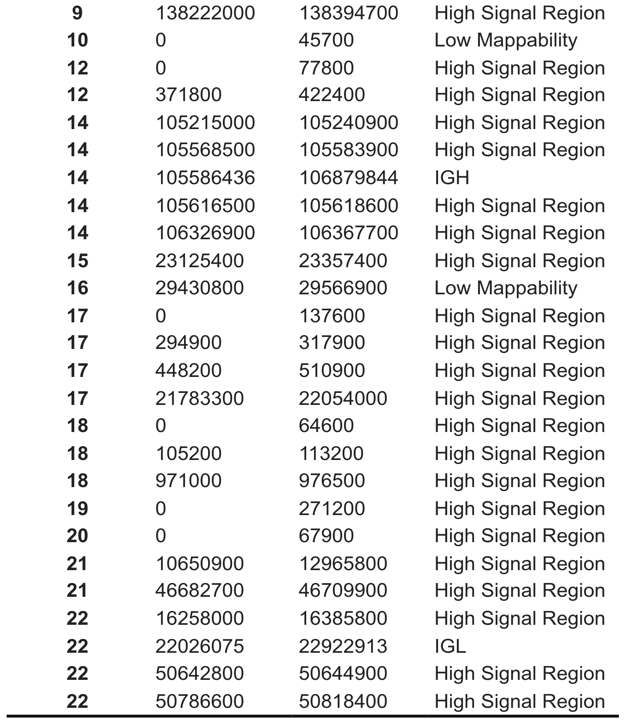

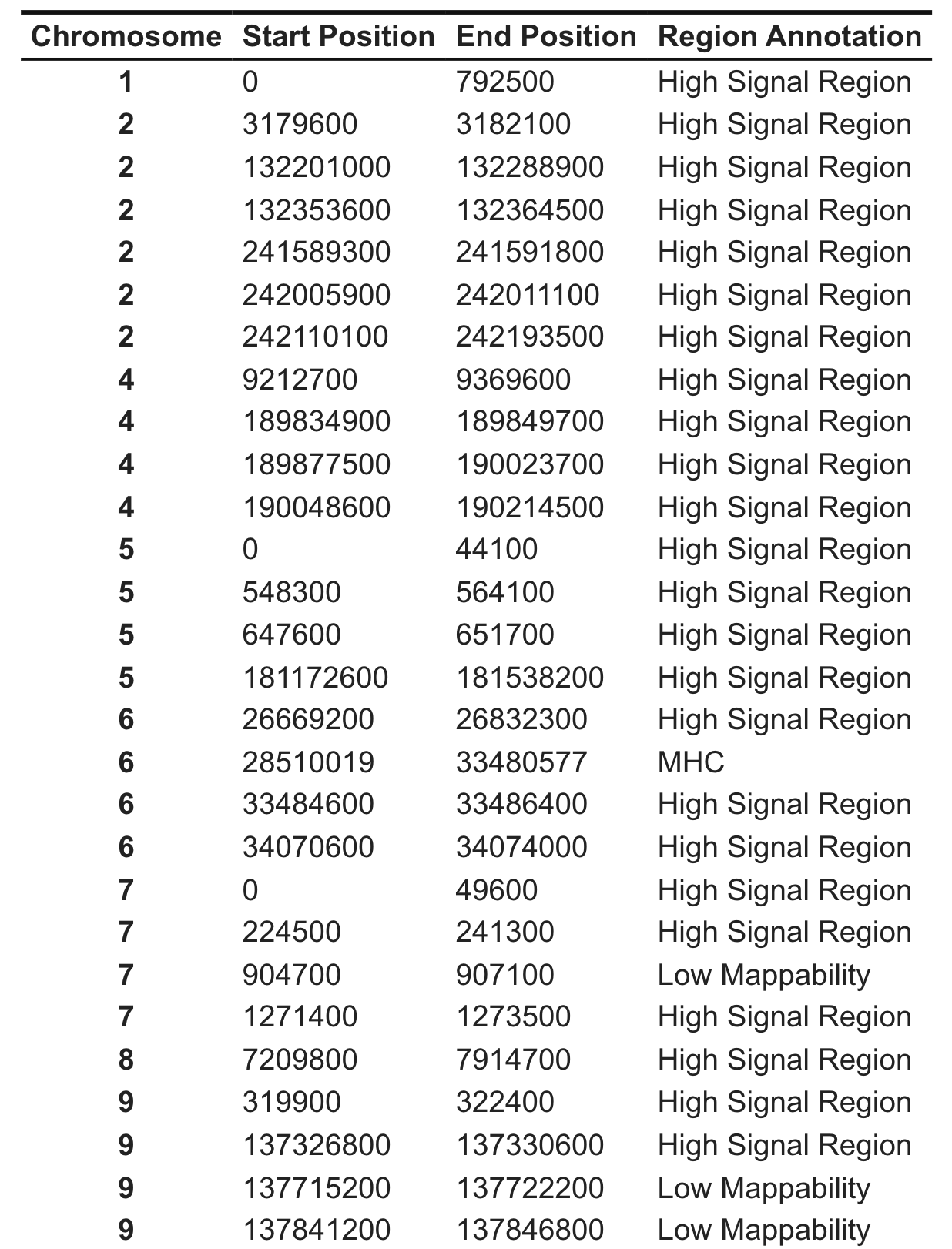

Unique genomic regions mapped to SNP-LAI pairs with low correlation in any of the three ancestries (African, Amerindian, and European), with start and end position and respective region annotations from either the ENCODE blacklist or UCSC genome browser gene cluster annotations. High signal region indicates potential unannotated repeats in the genome.

#### Supplementary Table 2: Effect size estimates comparison among the most significant variants from RFMix, FLARE3, and FLARE7 local ancestry inference

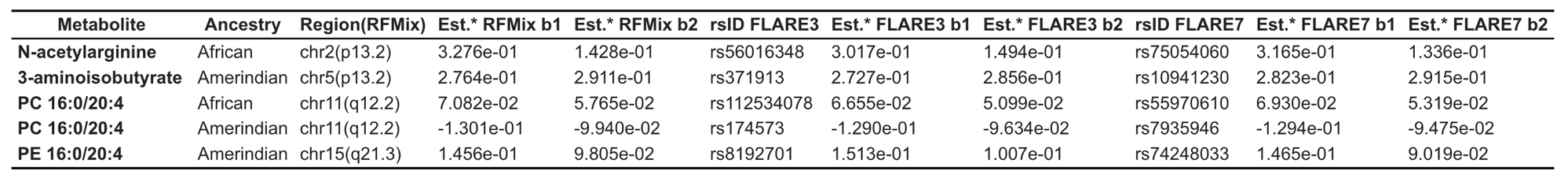

Comparison of effect size estimates (Est.*) from the most significant variants detected in the most significant genomic regions from RFMix admixture mapping results. The Est.* values are compared across the three inferences (RFMix, FLARE3, and FLARE7) as well as between the discovery batch 1 (b1) and replication batch 2 (b2).
